# mtTF1: A Novel Factor Involved in Mitochondrial Gene Expression in *Trypanosoma brucei*

**DOI:** 10.1101/2025.05.22.655275

**Authors:** Bianca Manuela Berger, Himaja Manjunatha, Markus Gerber, Torsten Ochsenreiter

**Author notes:** Corresponding author: Torsten Ochsenreiter, Institute of Cell Biology, University of Bern, Baltzerstrasse 4, Bern 3012.

## Abstract

Mitochondrial DNA replication and gene expression are essential for cell survival. In *Trypanosoma brucei*, a protozoan animal and human parasite, the mitochondrial RNA polymerase (mtRNAP) plays roles in both transcription and DNA replication. This study identifies and characterizes the first mitochondrial transcription factor (mtTF1) in the Kinetoplastea. mtRNAP and mtTF1 form a high-molecular-weight complex that localizes to the kinetoplast DNA (kDNA) and is essential for parasite survival in both life cycle stages. Their localization is interdependent, and both proteins influence maxicircle replication, but not minicircle replication. Knockdown of either protein result in altered gene expression, particularly affecting the minor strand of the mitochondrial genome. Since mtTF1 is unique to the Kinetoplastea, it might prove to be a promising drug target.

## Introduction

Mitochondria are membrane-bound organelles found in nearly all eukaryotic cells. Often referred to as the “powerhouse of the cell”, they generate energy in the form of ATP through oxidative phosphorylation, making them essential for cell survival and viability (Ernster & Schatz, 1981). Consistent with the endosymbiont theory, mitochondria possess their own DNA, reflecting their bacterial ancestry (Gray et al., 1999). However, over the course of evolution, the mitochondrial genome was strongly reduced, and some genes were transferred to the nucleus (Gray et al., 1999). Consequently, many mitochondrial proteins require import into the mitochondria (Pfanner et al., 2019). In contrast to nuclear gene expression, mitochondrial gene expression relies on a distinct set of proteins, most of which are encoded in the nucleus (Attardi, 1981; Gustafsson et al., 2016). According to the endosymbiont theory, a eubacterial-like multisubunit RNA polymerase would be expected in mitochondria. However, the mitochondrial RNA polymerase (mtRNAP, also known as POLRMT), the single subunit enzyme responsible for transcribing mitochondrial DNA (mtDNA) into RNA, shares structural and functional similarities with the T7 phage RNA polymerases and is conserved across most eukaryotes (Masters et al., 1987). While T7 phage RNA polymerases transcribe genes in a transcription factor-independent manner, all known eukaryotic mtRNAPs require transcription factors for their function (Cheetham et al., 1999; Jeruzalmi & Steitz, 1998). These transcription factors vary significantly across species. In human mitochondria, for instance, transcription involves four transcription factors (Gustafsson et al., 2016; Hillen et al., 2018). Two of which are involved in transcription initiation: mitochondrial transcription factor A (TFAM) binds and recruits mtRNAP to promoters, while transcription factor B2 (TFB2M) helps to open the DNA (Falkenberg et al., 2002; Hillen et al., 2017; Mangus et al., 1994). Another factor, mitochondrial transcription elongation factor (TEFM), replaces TFB2M and enhances the processivity of mtRNAP, thereby promoting the elongation of RNA transcripts (Hillen et al., 2017). Transcription initiated at the light strand promoter (LSP) is terminated by the mitochondrial transcription termination factor (mTERF), which acts as a roadblock (Agaronyan et al., 2015). In contrast to human mitochondrial transcription, yeast mitochondrial transcription requires only one mitochondrial transcription factor (Mtf1), which is homologous to TFB2M found in humans (De Wijngaert et al., 2021; Jang & Jaehning, 1991; Mangus et al., 1994; Paratkar & Patel, 2010). Transcription factors are much less conserved across eukaryotes compared to the mtRNAP, with considerable variation in their structures, regulatory mechanisms, and roles.

Besides mitochondrial transcription, mtRNAP is also involved in primer synthesis for mtDNA replication (Gustafsson et al., 2016). Interestingly, transcription from the light strand promoter (LSP) can be prematurely terminated, with the resulting short RNA acting as a primer for replication from the heavy strand origin of replication (OriH) (Falkenberg et al., 2007). Once the replication machinery completed two-thirds of the heavy-strand replication, the light-strand origin (OriL) becomes single-stranded and forms a stem-loop structure, which is accessible to mtRNAP. mtRNAP then synthesises the primer at OriL, which is subsequently used for light-strand replication (Clayton, 1991; Fusté et al., 2010; Robberson et al., 1972). In this strand-displacement model, which is the currently most-accepted model, no Okazaki fragments are required for lagging strand synthesis in humans (Yasukawa & Kang, 2018). Importantly, mtRNAP regulates the switch between primer formation and gene expression (Kühl et al., 2016). Knockout experiments revealed that under low mtRNAP levels, transcription at the LSP is favoured, ensuring primer formation and, consequently, a preference for replication over transcription (Gustafsson et al., 2016; Kühl et al., 2016).

*Trypanosoma brucei* (*T. brucei*) is the parasitic protozoan responsible for the diseases African trypanosomiasis (sleeping sickness) in humans and Nagana in cattle. Its metabolism strategies are tailored to different environments, allowing the parasite to grow in the mammalian host or the tsetse fly vector (Barrett et al., 2003; Malvy & Chappuis, 2011). These metabolic adaptations are essential for survival under diverse physiological conditions. In the mammalian bloodstream, the bloodstream form (BSF) trypanosomes rely almost exclusively on glycolysis for ATP production as glucose is abundant. In this stage, the single mitochondrion is reduced, and oxidative phosphorylation is downregulated (Besteiro et al., 2005). In contrast, in the tsetse fly vector, trypanosomes exist as procyclic form (PCF), which have fully developed and functional mitochondria that allow them to use oxidative phosphorylation for ATP generation via the electron transport chain (Besteiro et al., 2005; Matthews, 2005). The mechanisms underlying this metabolic plasticity are not yet fully understood. Hypotheses include the involvement of life cycle-specific transcription factors, post-transcriptional mechanisms such as editing or processing, as well as mRNA degradation (Aphasizheva et al., 2020).

In addition to the differential requirements for mitochondrial gene expression depending on the life cycle stage, an additional layer of complexity is added through the complex mitochondrial DNA network found in trypanosomes. The mitochondrial DNA is composed of two different types of circular DNA molecules (Shapiro et al., 1999). There are 25 maxicircles (23 kb in size), which encode proteins of the respiratory chain, two rRNAs, and two ribosomal proteins (Aphasizheva et al., 2020; Lukeš Jr et al., 2010). These maxicircles are similar to other eukaryotic mtDNA molecules, with the major difference being that no tRNAs are encoded in the mitochondrial DNA of trypanosomes, resulting in a strong dependence on tRNA import into the mitochondrion (Hancock & Hajduk, 1990; Simpson et al., 1989). In addition to maxicircles, trypanosomes contain approximately 5000 minicircles per cell, each around 1 kb in size. The minicircles encode for three to five gRNAs required for mRNA editing of maxicircle-encoded transcripts (Pollard et al., 1990; Sturm & Simpson, 1990). Interestingly, it has been shown that mtRNAP not only transcribes maxicircles, but that the protein is also involved in transcription of gRNAs encoded on minicircles (Grams et al., 2002; Hashimi et al., 2009). It is still unknown whether gene expression of these two DNA molecules is regulated by a common or different transcription factors.

While *T. brucei* strongly depends on the maintenance and expression of its mitochondrial DNA (kinetoplast DNA, kDNA), the closely related *T. evansi* and *T. equiperdum* can survive without a functional kinetoplast (Lai et al., 2008; Schnaufer et al., 2005). Importantly, these species are transmitted independently of the tsetse fly (Lai et al., 2008; Schnaufer et al., 2005). Survival without functional kDNA can be acquired through a single amino acid change in the nuclear-encoded y subunit of the catalytic F_1_ complex of the ATP synthase (Schnaufer et al., 2005). Similarly, the mutation of a single leucin at position 262 to proline (yL262P) allows the yL262P BSF trypanosome to survive without functional mitochondrial gene expression and/or replication. (Dean et al., 2013; Schnaufer et al., 2005). When mitochondrial DNA replication is impaired, the yL262P cells lose their kDNA without affecting cell survival. These findings suggest that the BSF trypanosomes only rely on the expression of the A6 subunit of the ATP synthase (and the rRNAs and ribosomal proteins required for its translation). This dependence can be overcome by the yL262P mutation, rendering the cells independent of mitochondrial gene expression (Dewar et al., 2022; Schnaufer et al., 2005).

In the present study, immunoprecipitation experiments of the mtRNAP led to the identification of the first mitochondrial transcription factor (mtTF1) in *T. brucei.* Further experiments confirmed the co-localization of mtRNAP and mtTF1 at the kDNA and demonstrate their interdependence. We show that mtTF1 is the first transcription factor identified in the Kinetoplastea with involvement in both maxicircle and minicircle transcription. Upon mtTF1 knockdown, we observed strand-specific effects on maxicircle transcription, with the minor strand transcripts showing a more significant decrease compared to major strand transcripts. A similar observation was made upon ablation of mtRNAP. It remains to be elucidated whether the promoters for major and minor strand transcription have different strengths, accounting for this phenotype. Additionally, both mtRNAP and mtTF1 are required for maxicircle but not minicircle replication. Interestingly, mtTF1 is conserved among all Kinetoplastea, including human pathogens such as *Leishmania* and *Trypanosoma cruzi*. Given that mtTF1 is essential in *T. brucei*, it may represent an interesting candidate for drug development.

## Results

### The mitochondrial RNA-polymerase (mtRNAP) localizes to the kDNA and is essential in the procyclic and bloodstream form trypanosomes

The mitochondrial RNA-polymerase (mtRNAP) of *Trypanosoma brucei (T. brucei)* has been identified by Grams and coworkers in 2002 (Grams et al., 2002). The authors showed the involvement of mtRNAP in maxicircle replication, maxicircle transcription and the essentiality of the protein for the procyclic form (PCF) of *T. brucei* (Grams et al., 2002). Additionally, Hashimi and coworkers could show that mtRNAP is also involved in minicircle transcription (Hashimi et al., 2009). To further investigate the role of mtRNAP, we first C-terminally tagged mtRNAP with a 3xMyc tag and determined the localization of the protein. Biochemical analysis using digitonin fractionation showed that mtRNAP-Myc localizes to the mitochondria-enriched fraction (Figure 1A). The tagged mtRNAP-Myc is found in the mitochondria-enriched pellet, whereas no signal was detected in the supernatant containing the cytosolic proteins (Figure 1A). With immunofluorescence analysis (IFA), we observed mtRNAP-Myc localization at the kDNA throughout the cell cycle in the PCF (Figure 1B). Next, inducible mtRNAP RNAi cell lines were generated in PCF and bloodstream form (BSF) trypanosomes. The RNAi effectively resulted in depletion of the protein as shown by Western blot (Figure 1C and D inlets). RNAi depletion of mtRNAP results in a growth retardation starting after two to three days of induction in the PCF (Figure 1C). Since the PCF and BSF have differential dependence on mitochondrial oxidative phosphorylation for ATP production, we wanted to investigate if mtRNAP is essential in the BSF as well. Depletion of mtRNAP in BSF results in a growth defect starting after two to three days of induction, indicating that mtRNAP is essential in both the PCF and BSF of *T. brucei*.

**Figure 1:**
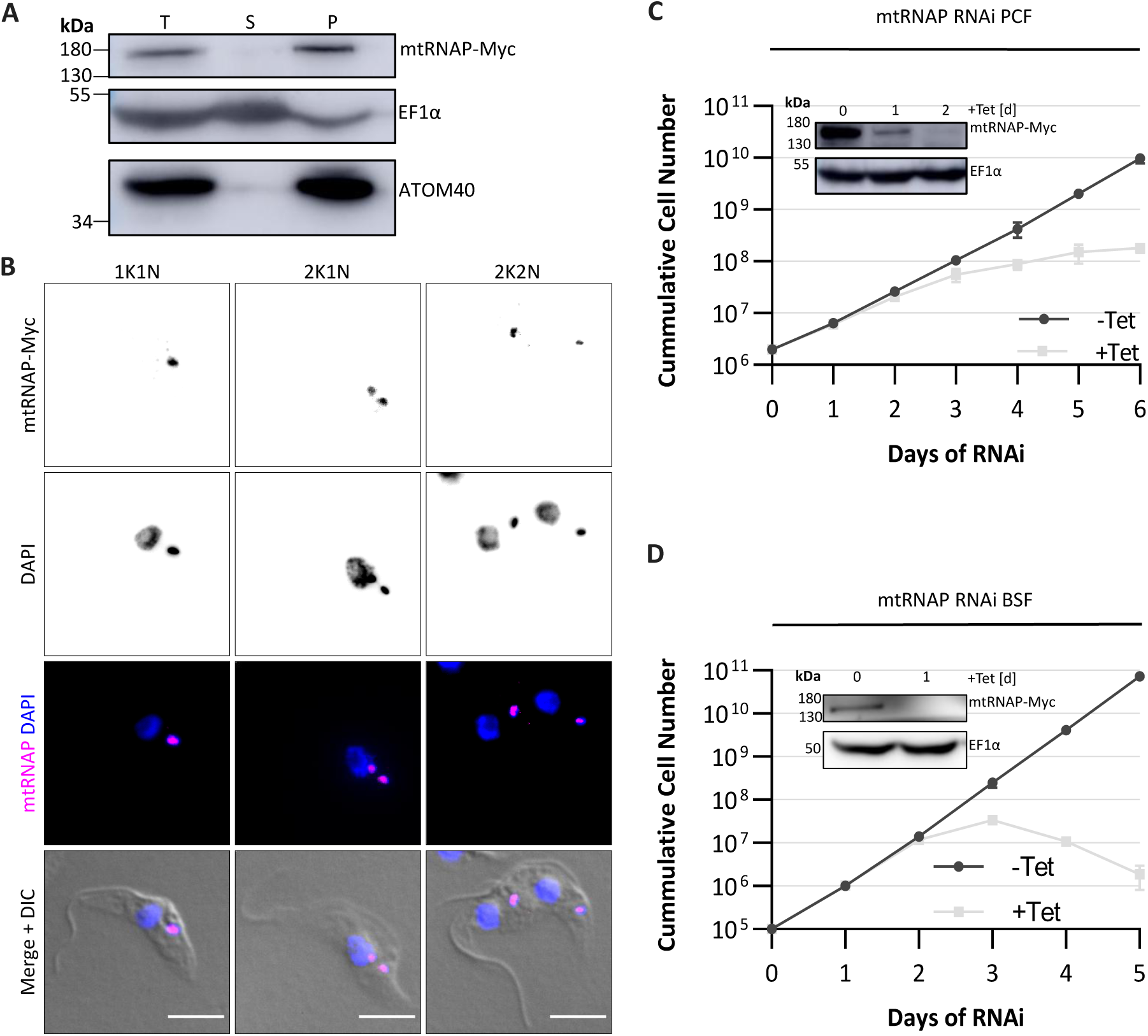
mtRNAP localizes to the kDNA and is essential in the procyclic form (PCF) and bloodstream form (BSF) of trypanosomes. **(A)** Western blot of total cell lysate (T), the soluble cytosolic fraction (S) and the digitonin-extracted mitochondria-enriched pellet (P) of cells expressing C-terminally Myc-tagged mtRNAP. The Western blot was probed for EF1α and ATOM40, which serve as cytosolic and mitochondrial markers, respectively. **(B)** Immunofluorescence images showing localization of the Myc-tagged mtRNAP (magenta) at the kDNA throughout the cell cycle (K = kDNA, N = nucleus). DNA is stained with DAPI (blue). Cells are visualized in DIC (differential interference contrast microscopy). **(C and D)** Growth curve of non-induced (black, -Tet) and mtRNAP RNAi-induced (grey, +Tet) cells in the procyclic form (PCF) and bloodstream form (BSF), respectively. Inset: Western blot of non-induced and mtRNAP RNAi-induced cells (+Tet). Western blot was probed for Myc to detect mtRNAP levels and EF1α, which served as a loading control. Scale bars in B = 5 µm. Data are shown as mean +/-standard deviation (SD). n = 3 for C and D.

### Immunoprecipitation of mtRNAP to identify interaction partners of mtRNAP in *T. brucei*

To identify interaction partners of mtRNAP in *T. brucei*, we performed immunoprecipitation with mtRNAP tagged C-terminally with an HA tag (PCF) or Myc tag (BSF). For the immunoprecipitation, cells were lysed, and mitochondria were enriched using digitonin fractionation. The workflow is shown in Supplementary Figure 1. The input, flowthrough and immunoprecipitation fraction were analysed by Western blot (Figure 2A). mtRNAP-HA is detected in the input, as well as the immunoprecipitation fraction, whereas the outer mitochondrial membrane protein ATOM40 (archaic translocase of the mitochondrial outer membrane 40) is found in the input and flowthrough (Figure 2A). Mass spectrometry revealed that mtRNAP and the hypothetical protein Tb927.6.4510 (here after called mitochondrial transcription factor 1: mtTF1) were the most enriched proteins in the mtRNAP pulldown experiments in PCF as well as BSF trypanosomes (Figure 2B and C). Table 1 and 2 summarize the significantly enriched proteins in the mtRNAP immunoprecipitation performed in PCF and BSF, respectively, with a fold changes larger than two.

**Figure 2:**
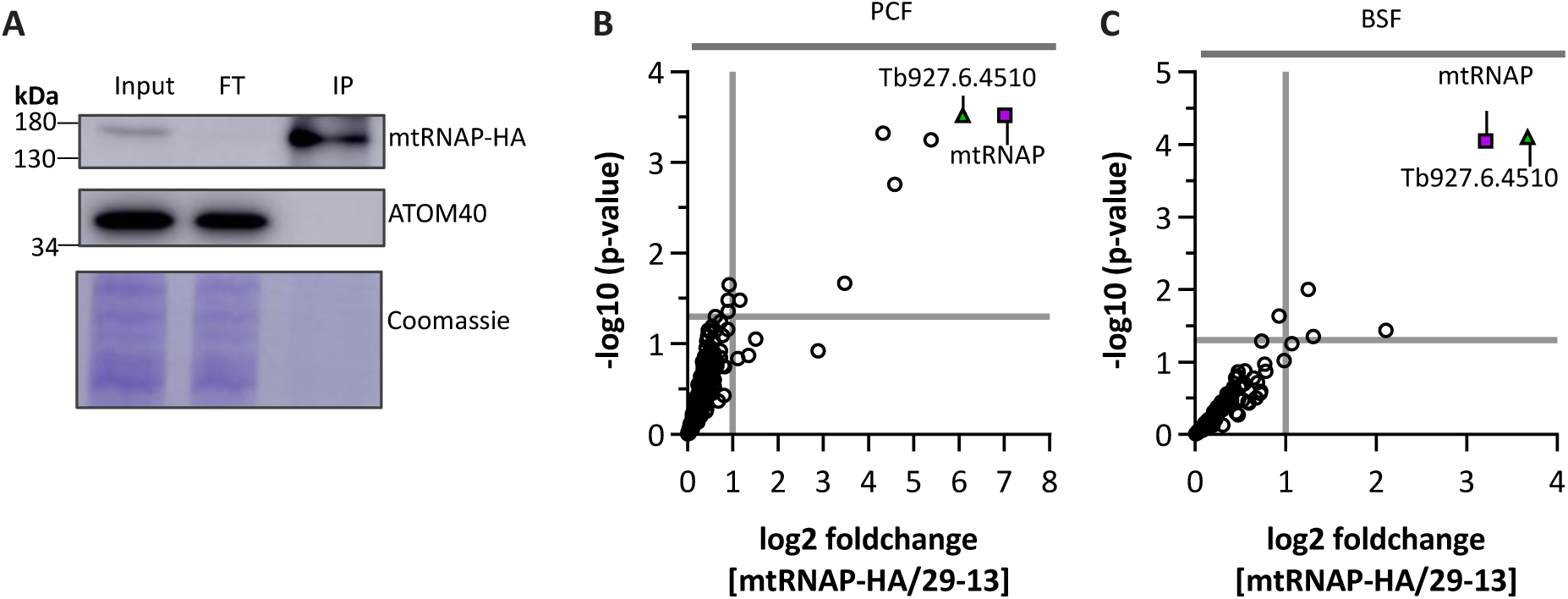
Immunoprecipitation analysis of mtRNAP to identify interaction partners. **(A)** Western blot analysis of the immunoprecipitation performed with PCF cells expressing HA-tagged mtRNAP. Lysed mitochondria were used as input for the immunoprecipitation. The mitochondrial protein ATOM40 serves as a control and is expected in the flowthrough (FT), while mtRNAP is expected in the immunoprecipitation fraction (IP). **(B and C)** Volcano plot of the immunoprecipitation of cells expressing the tagged mtRNAP in the procyclic form (PCF) and the bloodstream form (BSF), respectively. Wild-type cells without tagged mtRNAP served as negative control for background subtraction. Horizontal lines indicate a p-value of 0.05 and vertical lines mark a two-fold enrichment of proteins in the tagged cell lines compared to wildtype cells. n = 3 for B and C.

**Table 1:** Significantly enriched proteins in the mtRNAP immunoprecipitation in the PCF. Shown are hits with a fold change larger than two.

| Rank of log2 FC | Accession Number | Protein | Location based on TrypTag (C-terminal tagging) |
| --- | --- | --- | --- |
| 1 | Tb927.11.5780 | mtRNAP | Cytoplasm, mitochondrion |
| 2 | Tb927.6.4510 | hypothetical protein (mtTF1) | Kinetoplast |
| 3 | Tb927.9.7670 | hypothetical protein | Nucleolus |
| 4 | Tb927.11.15850 | mitoribosomal SSU assembly factor 5 | Cytoplasm, mitochondrion |
| 5 | Tb927.11.6980 | phosphoenolpyruvate carboxykinase | Mitochondrion, kinetoplast |
| 6 | Tb927.9.1980 | hypothetical protein | Axoneme, cytoplasm |
| 7 | Tb927.11.3120 | nucleolar GTP-binding protein 1 | Nucleolus |

**Table 2:** Significantly enriched proteins in the mtRNAP immunoprecipitation in the BSF. Shown are hits with a fold change larger than two.

| Rank of log2 FC | Accession Number | Protein | Location based on TrypTag (C-terminal tagging) |
| --- | --- | --- | --- |
| 1 | Tb927.6.4510 | hypothetical protein (mtTF1) | Kinetoplast |
| 2 | Tb927.11.5780 | mtRNAP | Cytoplasm, mitochondrion |
| 3 | Tb927.9.4210 | fatty acyl CoA synthetase 3 | - |
| 4 | Tb927.10.5770 | valosin-containing protein | Nucleoplasm, cytoplasm |
| 5 | Tb927.5.1300 | V-type proton ATPase subunit a | Endocytic |

### mtTF1 is conserved in the Kinetoplastea and co-localizes with mtRNAP at the kDNA

mtTF1 is a 37 kDa protein with an isoelectric point (IP) of 6.9, which is in a similar range to the IP of mtRNAP (IP = 7.07). mtTF1 is conserved in the Kinetoplastea but not found in other eukaryotic organisms (Figure 3A). AlphaFold3 was used to predict the structure of mtTF1 (Figure 3B) (Abramson et al., 2024). The N-terminal domain of the mtTF1 prediction has a lower confidence score and potentially contains three alpha helices. The middle part of the predicted protein structure until the C-terminus, is composed of helix-turn-helix repeats, predicted with high confidence (Figure 3B). To verify the interaction between mtRNAP and mtTF1, a double-tagged cell line was generated in the PCF: mtRNAP-HA and mtTF1-Myc. Co-immunoprecipitation with beads coupled to anti-HA antibodies (pulldown of mtRNAP-HA) showed that mtRNAP-HA as well as mtTF1-Myc were in the co-immunoprecipitation fraction (Figure 3C). Similar results were obtained performing reciprocal co-immunoprecipitation with beads coupled to anti-Myc antibodies (pulldown of mtTF1-Myc) (Figure 3D). Importantly, samples for co-immunoprecipitation were treated with RNase and DNase, indicating that the interaction between mtRNAP and mtTF1 is RNA- and DNA-independent (Figure 3C and D). IFA of mtRNAP and mtTF1 showed that the two epitope-tagged proteins colocalize at the kDNA throughout the cell cycle (Figure 3E) in the PCF. For better resolution of mtRNAP and mtTF1 and to calculate the Pearson correlation coefficient, we performed Expansion Microscopy (ExM) with the Pre-Staining Expansion Microscopy (PS-ExM) protocol developed by Atchou and coworkers (Atchou et al., 2023). Signal of mtRNAP and mtTF1 overlap at the kDNA disk in the PCF with a Pearson correlation coefficient of 0.80 ± 0.07 (the average ± SEM of ten kDNAs was calculated with Fisher’s transformation).

**Figure 3:**
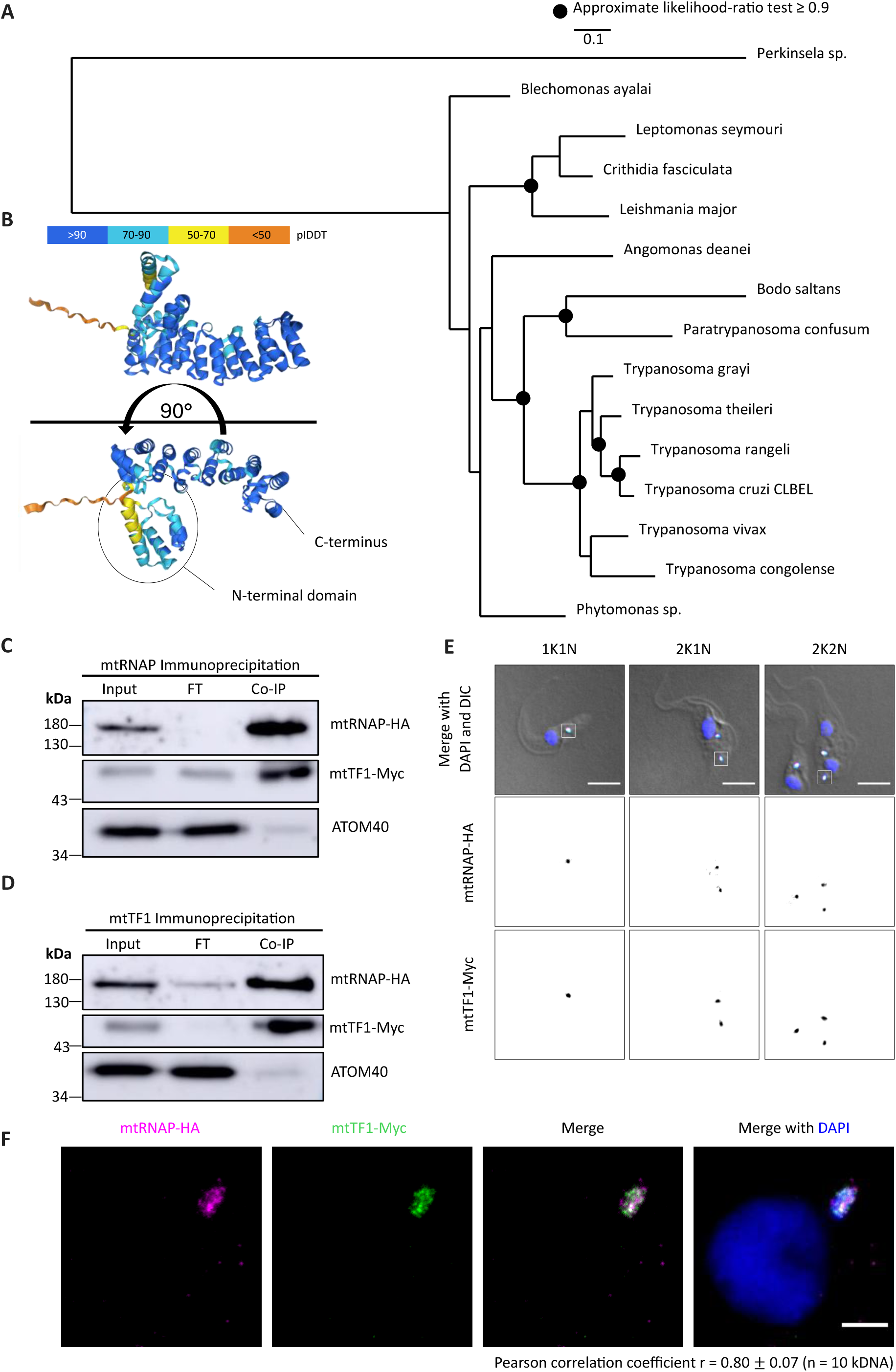
mtTF1 is conserved in the Kinetoplastea and co-localizes with mtRNAP at the kDNA in the PCF. **(A)** Phylogenetic tree illustrating the conservation of mtTF1 among the Kinetoplastea. **(B)** AlphaFold3 prediction of mtTF1. The N-terminal domain is marked with a circle. **(C and D)** Western blot of the co-immunoprecipitation (Co-IP) performed with cells expressing the HA-tagged mtRNAP and Myc-tagged mtTF1 using anti-HA and anti-Myc beads, respectively. Digitonin-extracted mitochondria were used as input for the Co-IP. Western blots were probed with anti-HA, anti-Myc and anti-ATOM40 antibodies. ATOM40 serves as negative control, which is not expected in the Co-IP fraction, but only in the flowthrough (FT). **(E)** Immunofluorescence images showing the localization of mtRNAP-HA (magenta) and mtTF1-Myc (green) throughout the cell cycle (K = kDNA, N = nucleus). DNA is visualized with DAPI (blue), and cells are visualized in DIC (differential interference contrast microscopy). **(F)** Expansion microscopy images of cells expressing mtRNAP-HA (magenta) and mtTF1-Myc (green). DNA is stained with DAPI (blue). Pearson correlation coefficient in panel F was calculated for n = 10 kDNA. Data are presented as mean +/-SEM. Scale bars = 5 µm.

### mtTF1 is essential in the PCF and its localization depends on mtRNAP

Conditional knockdown of mtTF1 was performed by inducible RNAi, which results in depletion of the protein as verified with Western blot using the mitochondrial heat shock protein 70 (mtHSP70) as a loading control (Figure 4A). Interestingly, we observed a decrease in protein levels of the nuclear-encoded cytochrome c oxidase subunit 4 (COX4) after four days of mtTF1 RNAi induction, indicating that mtTF1 is important for maintenance of respiratory chain complexes. Growth retardation is observed starting after three to four days of mtTF1 RNAi induction (Figure 4B). To test if the proper localization of mtTF1 at the kDNA depends on the presence of mtRNAP, we generated a cell line with an HA-epitope tagged mtTF1 (C-terminal tagging) and inducible mtRNAP RNAi. The growth curve is shown in Supplementary Figure 2A. Two days after induction of mtRNAP knockdown, mtTF1 did not localize properly to the kDNA but was dispersed within the mitochondrion as assessed by IFA (Figure 4C). These findings were quantified by measuring the signal of HA-tagged mtTF1 at the kDNA and next to the kDNA in control (-Tet) and mtRNAP depleted cells (+Tet [2d]) (Figure 4D). While the signal intensity of mtTF1-HA at the kDNA decreased upon mtRNAP RNAi, the signal intensity of mtTF1-HA next to the kDNA increased (Figure 4D). To verify if mtTF1 still localizes to the mitochondrion upon mtRNAP depletion, we performed digitonin extraction. mtTF1-HA was detected in the mitochondria-enriched fraction in the control cells and upon mtRNAP depletion (Figure 4E). Blue-Native PAGE confirmed that mtRNAP-HA and mtTF1-HA are in a high-molecular weight complex of similar size (Supplementary Figure 2B). Next, we tested the integrity of the high-molecular weight complex upon knockdown of mtRNAP. In the tested condition, mtRNAP depletion leads to loss of the high-molecular weight complex containing mtTF1-HA (Figure 4F). Western blot of a duplicate experiment revealed no effect on mtTF1-HA levels upon mtRNAP RNAi two days after induction (Figure 4F). Thus, we conclude that (i) the localization of mtTF1 to the kDNA depends on mtRNAP and (ii) the formation of the mtTF1-containing high-molecular weight complex requires the presence of the mtRNAP.

**Figure 4:**
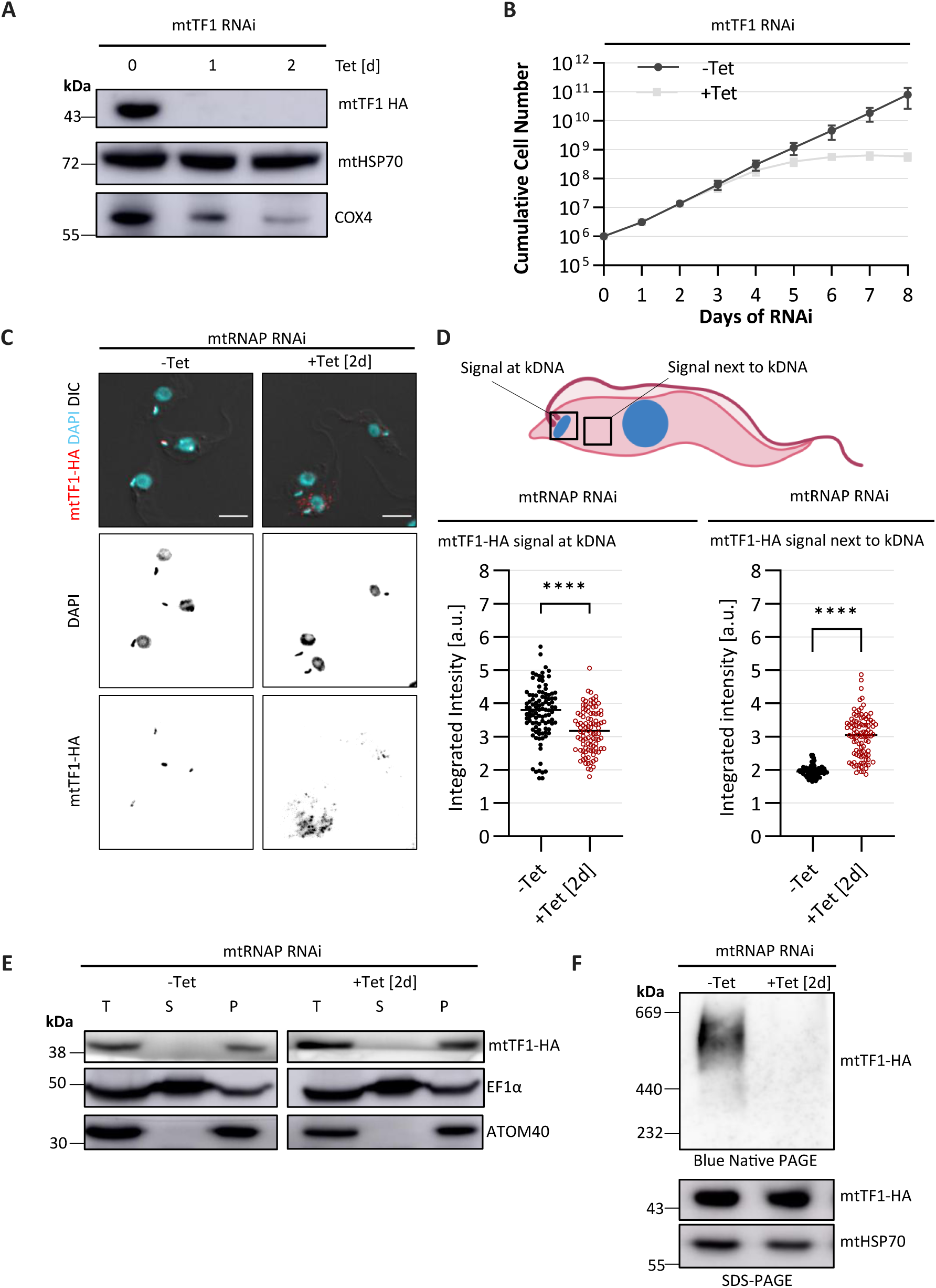
**mtTF1 is essential in the insect form, and its localization at the kDNA depends on mtRNAP**. **(A)** Western blot analysis showing depletion of mtTF1 after one and two days of tetracycline-induced mtTF1 RNAi compared to non-induced control cells. The Western blot was probed for COX4 to assess the integrity of mitochondrial respiratory chain complexes, while mtHSP70 serves as a loading control. **(B)** Growth curve of PCF trypanosomes, upon tetracycline-induced mtTF1 RNAi (+Tet) cells compared to non-induced control cells (-Tet). **(C)** Immunofluorescence images showing mtTF1-HA localization (red) in control cells (-Tet) and upon induction of mtRNAP RNAi (+Tet[2d]). DNA is stained with DAPI (cyan). Cells are visualized with DIC (differential interference contrast microscopy). **(D)** Quantification of mtTF1-HA signal intensity at the kDNA (left) and next to the kDNA (right) upon mtRNAP RNAi (+Tet [2d], red) compared to non-induced control cells (-Tet, black). **(E)** Digitonin extraction was performed in control cells (-Tet) and mtRNAP RNAi-induced cells (+Tet [2d]) to biochemically investigate the localization of mtTF1 upon mtRNAP depletion. Western blot analysis of total cell lysate (T), the soluble cytosolic fraction (S) and digitonin-extracted mitochondria-enriched pellet (P). The Western blot was probed for HA to detect mtTF1. EF1α and ATOM40 serve as cytosolic and mitochondrial marker, respectively. (F) Blue Native PAGE analysis (top) of mtTF1-HA in mtRNAP RNAi-induced cells (+Tet [2d]) and non-induced controls (-Tet). Western blot (bottom) probed for HA to detect mtTF1 upon mtRNAP depletion (+Tet [2d]) compared to non-induced controls (-Tet). Blue Native PAGE and SDS-PAGE were performed as duplicate experiments. Scale bars = 5 µm. Data are shown as mean +/-standard deviation (SD). n = 3 for B and F. n = 100 for E.

### mtRNAP localization to the kDNA depends on mtTF1

Knockdown of mtTF1 was used to investigate how the localization of mtRNAP and the high-molecular weight complex containing mtRNAP depend on the novel identified factor. The growth curve is shown in Supplementary Figure 2C. When mtTF1 is depleted, mtRNAP intensity at the kDNA is reduced (Figure 5A and B), while the signal of mtRNAP next to the kDNA is not affected by mtTF1 RNAi (Figure 5C). Next, we performed Blue Native PAGE to test whether mtTF1 depletion affects the high-molecular weight complex. We observed a decrease in the size of the complex when mtTF1 is depleted, indicating loss of mtTF1 from the complex (Figure 5D and Supplementary Figure 2D). mtRNAP-HA levels were not affected by mtTF1 knockdown as assessed by Western blot analysis (Figure 5E). These results show that mtRNAP is still present in a high-molecular weight complex upon mtTF1 RNAi. Similar to the observation that mtTF1 localization depends on mtRNAP, the localization of mtRNAP at the kDNA strongly depends on mtTF1.

**Figure 5:**
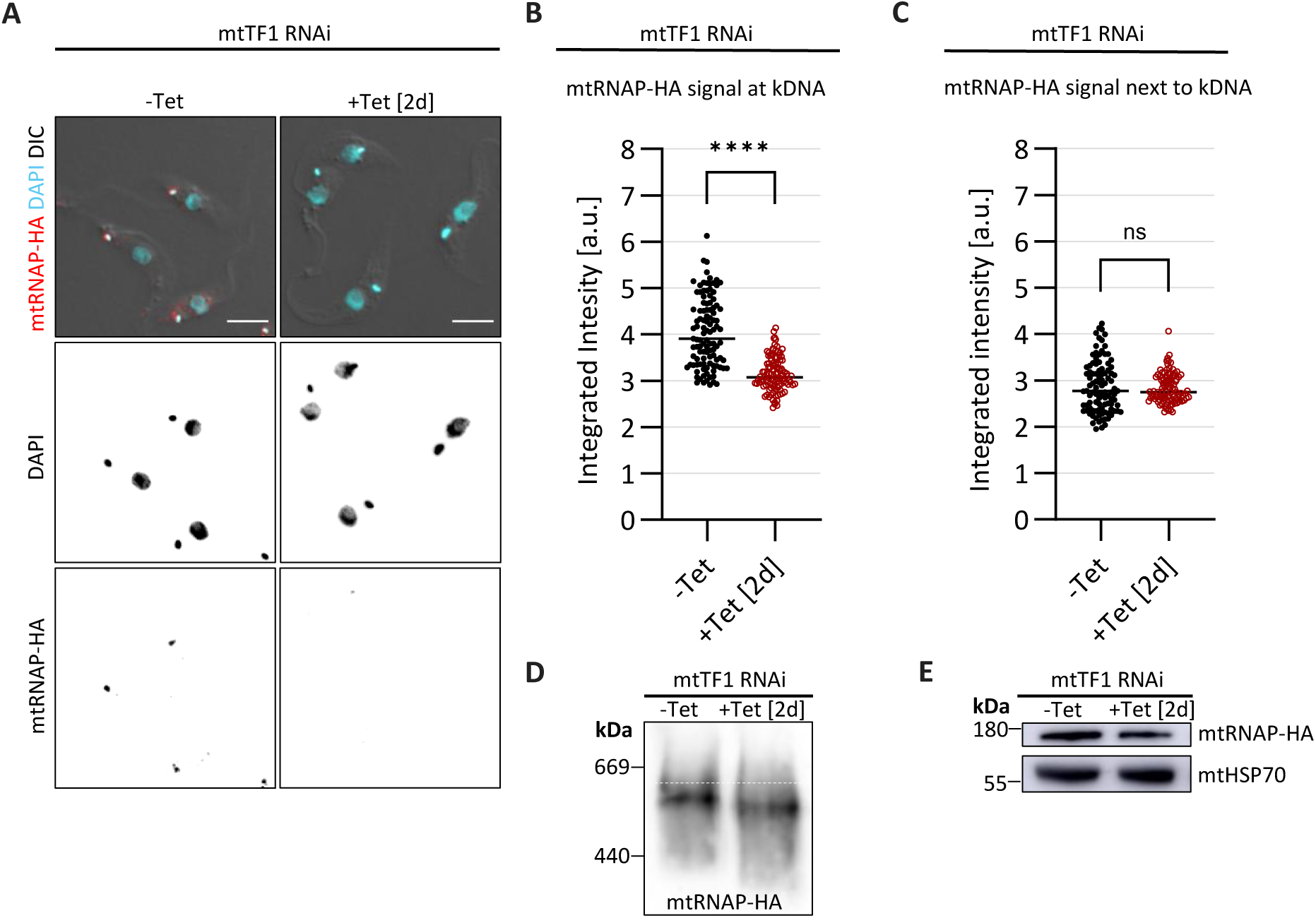
mtRNAP intensity at the kDNA decreases upon mtTF1 RNAi in the PCF. **(A)** Immunofluorescence images showing mtRNAP-HA localization (red) upon mtTF1 RNAi. DNA is stained with DAPI (cyan). Cells are visualized with DIC (differential interference contrast microscopy). **(B and C)** Quantification of the mtRNAP-HA signal intensity assessed by immunofluorescence imaging upon mtTF1 knockdown (+Tet [2d], red) compared to non-induced control cells (-Tet, black). mtRNAP-HA signal was measured at the kDNA and next to the kDNA, respectively. **(D)** Blue Native PAGE analysis probing for Ha-tagged mtRNAP in non-induced control cells (-Tet) and mtTF1 RNAi-induced cells (+Tet [2d]). **(E)** Western blot analysis of mtRNAP-HA levels in non-induced control cells (-Tet) and mtTF1 RNAi-induced cells (+Tet [2d]). The Western blot was probed against HA to detect mtRNAP and against mtHSP70, which serves as a loading control. Scale bars = 5µm. Data are shown as mean +/- standard deviation (SD). n = 3 for A, D and E. n = 100 for B and C.

### mtRNAP and mtTF1 are involved in maxicircle replication and transcription of mini- and maxicircles

The function of mtRNAP and mtTF1 in kDNA replication and maxicircle transcription were investigated with qPCR and RT-qPCR, respectively. Similarly to what was previously published by Grams and coworkers, we observed a decrease of maxicircle abundance but not minicircle DNA abundance two days after mtRNAP RNAi induction (Grams et al., 2002). Maxicircle DNA (quantified with primers amplifying ND4 and 12S rDNA) was reduced below 60% upon mtRNAP RNAi (Figure 6A). RNA sequencing was performed to obtain an overview of mini- and maxicircle transcript level changes following mtRNAP and mtTF1 knockdown. The results are shown in Supplementary Figure 3. We observed differential effects on individual mitochondrial transcripts. To further explore this observation, we performed RT-qPCR in biological triplicates on never-edited transcripts and found strand-dependent differences in maxicircle transcript levels (Figure 6B). While transcripts from the major strand (12S rRNA and ND4) went down to roughly 60-80% two days after induction of mtRNAP RNAi, the minor strand transcripts (COI and ND1) were more affected by mtRNAP knockdown with less than 30% detected in induced cells compared to control cells (Figure 6B). With Northern blot probing for the ND7 gRNA, we could show that mtRNAP is involved in gRNA synthesis which is in line with results from Hashimi and coworkers (Hashimi et al., 2009). This probe was chosen based on its high abundance in RNA sequencing experiments of control cells compared to mtRNAP depleted cells (+Tet [2d]).

**Figure 6:**
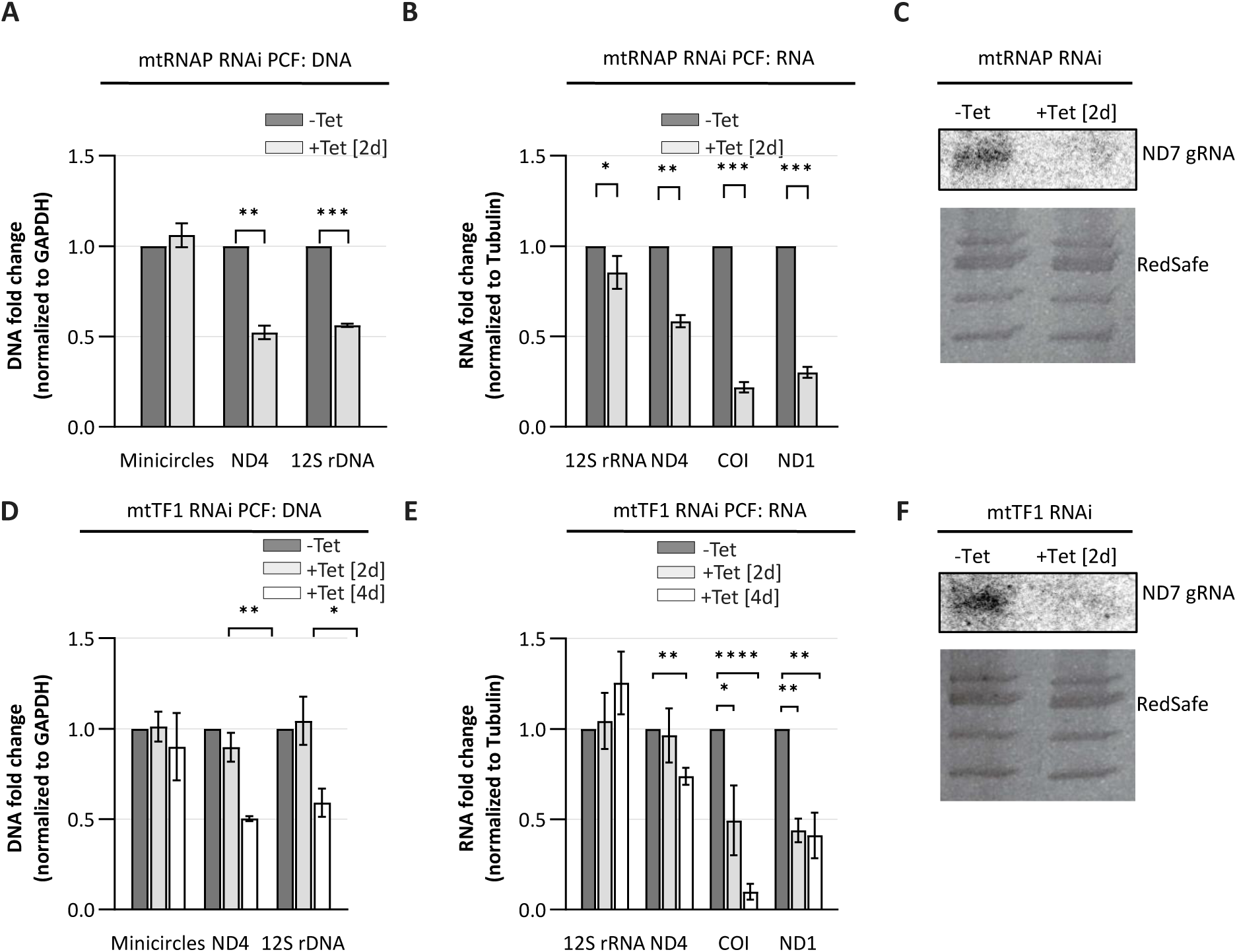
mtRNAP and mtTF1 are involved in maxicircle replication as well as maxi- and minicircle transcription in the PCF. **(A and B)** qPCR and RT-qPCR analysis of mini- and maxicircle DNA in control (-Tet) and mtRNAP RNAi-induced cells (+Tet [2d]). **(C)** Northern blot analysis probing for ND7 gRNA in control (-Tet) and mtRNAP RNAi-induced cells (+Tet [2d]). RedSafe-stained gels serve as loading control. **(D and E)** qPCR and RT-qPCR analysis in control (-Tet, dark grey) and mtTF1 RNAi-induced cells (+Tet [2d] in light grey and +Tet [4d] in white). **(F)** Northern blot probing for ND7 gRNA in control (-Tet) and mtTF1 RNAi-induced cells (+Tet [2d]). RedSafe-stained gels serve as loading control. n = 3 for graphs A, B, C, D, E and F. Data are shown as mean +/- standard deviation (SD).

To exclude the possibility that the effects on the major strand upon mtRNAP RNAi were simply because of the decreased maxicircle DNA abundance, we compared the mtRNAP RNAi data with knockdown of the other primase that has been shown to result in decreased maxicircle DNA abundance, which is Primase 1 (Pri1). Pri1 was described by Hines and Ray in 2010 (Hines & Ray, 2010). In contrast to mtRNAP, which localizes to the kDNA, Pri1 shows localization at the antipodal site (APS) throughout the cell cycle in the PCF (Supplementary Figure 4A). RNAi of Pri1 results in efficient depletion of the protein as determined by Western Blot and a growth retardation starting three to four days after RNAi induction (Supplementary Figure 4B). Interestingly, upon Pri1 RNAi, we observed fragmentation of the kDNA two days after RNAi induction (Supplementary Figure 4C, arrows) and subsequent shrinkage of the kDNA (Supplementary Figure 4C and D). Similarly to the results of Hines and Ray, qPCR confirmed a loss of maxicircle DNA below 50% two days after Pri1 RNAi induction. In contrast to the mtRNAP RNAi, Pri1 RNAi also affects minicircles that were reduced below 60% (Supplementary Figure 4E). Importantly, RT-qPCR did not show any effects on major or minor strand maxicircle transcripts. Therefore, we conclude that (i) the reduction of maxicircle DNA upon mtRNAP RNAi does not directly result in decrease of the maxicircle transcripts (ii) mtRNAP can compensate for the 50% loss of maxicircle DNA upon Pri1 RNAi, still synthesising normal transcript levels of maxicircles.

Next, we performed qPCR and RT-qPCR to determine if mtTF1 is involved in replication and/or transcription of the kDNA. Two days after mtTF1 RNAi induction, we did not observe changes in mtDNA abundance (Figure 6D). However, four days after RNAi induction, a time point just at the onset of the growth phenotype, the maxicircle DNA abundance decreased to 50%. Furthermore, two days after mtTF1 RNAi induction, we observed a decrease of transcripts encoded on the minor strand (COI and NDI), while major strand transcripts (12S rRNA and ND4) are not affected (Figure 6E). Interestingly, after two more days of mtTF1 knockdown, the major strand seemed to be affected as well with ND4 transcript levels decreasing roughly to 60%. The effects observed for mtTF1 RNAi are very similar to the knockdown of mtRNAP with an effect on maxicircle replication and transcription with strand-specific differences. Major strand transcription seems to be affected less by RNAi when compared to the minor strand. Northern blot with probes against ND7 gRNA further showed that mtTF1 is involved in minicircle transcription, as ND7 gRNA levels were reduced upon mtTF1 RNAi (+Tet [2d]) (Figure 6F).

### mtRNAP is dispensable in yL262P BSF

To further test the function of mtRNAP in *T. brucei*, we performed mtRNAP RNAi in the yL262P cell line. The yL262P cell line can grow under glycolytic conditions without a mitochondrial genome. Therefore, this cell line is a suitable tool to study the function of mtRNAP after a longer period of RNAi induction (which would be lethal for regular BSF or PCF trypanosomes). We performed mtRNAP RNAi and did not observe a growth phenotype upon induction, while we could confirm efficient RNAi induction by Western blot (Figure 7A). Interestingly, even after ten days of RNAi induction, no apparent phenotype was visible for the kDNA as observed with DAPI staining (Figure 7B). The kDNA size was measured in control cells and cells which have been induced for mtRNAP RNAi for ten days. This quantification did not reveal any changes (Figure 7C). With qPCR we saw a decrease of maxicircle DNA abundance below 50%, while minicircles were not affected (Figure 7D). The strand-specific effects that we observed with RT-qPCR in the PCF could be replicated with the knockdown of mtRNAP in yL262P. Upon mtRNAP RNAi in yL262P, the minor strand maxicircle transcript abundance decreased to less than 10% compared to non-induced cells (Figure 7E). While ND4 showed a decrease in mRNA abundance, the other tested major strand transcript (12S rRNA) showed a two-fold increase upon mtRNAP RNAi.

**Figure 7:**
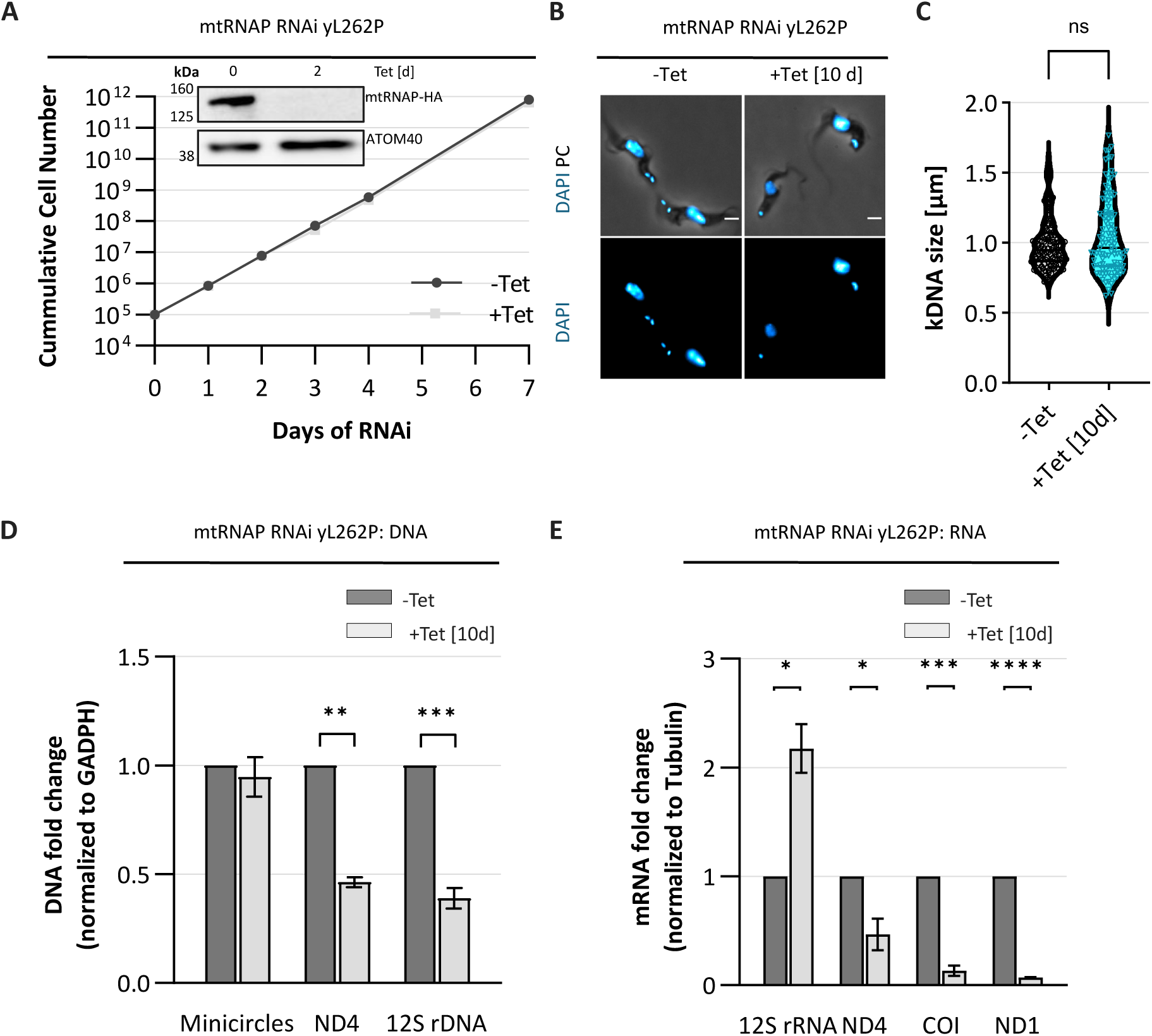
mtRNAP is dispensable in yL262P BSF trypanosomes. **(A)** Growth curve and Western blot analysis of non-induced (-Tet) and mtRNAP RNAi-induced (+Tet) yL262P cells. mtRNAP is C-terminally tagged with an HA tag in the yL262P cells and was detected with anti-HA antibodies. ATOM40 serves as a loading control. **(B)** DAPI staining of non-induced and mtRNAP RNAi-induced yL262P cells (+Tet [10d]). Cells are visualized with phase contrast (PH). **(C)** Quantification of kDNA size in non-induced (-Tet, black) and mtRNAP RNAi-induced yL262P cells (+Tet [10d], cyan). **(D and E)** qPCR and RT-qPCR analysis for control (-Tet, dark grey) and mtRNAP RNAi-induced (+Tet [10d], light grey) yL262P cells. Scale bars = 2 µm. n = 3 for panels A, D and E and n = 100 for C. Data are shown as mean +/- standard deviation (SD).

### mtRNAP and mtTF1 overexpression does not impair cell growth and does not alter mitochondrial gene expression

Since RNAi of mtRNAP and mtTF1 resulted in impaired cell growth, we wanted to examine if the overexpression of either mtRNAP or mtTF1 influences cell growth and mitochondrial gene expression. The tetracycline-inducible overexpression of mtRNAP tagged with a 3xHA revealed that the overexpressed protein localizes not only at the kDNA but throughout the mitochondrial organelle (Supplementary Figure 5A). We also found that overexpression does not impair cell growth (Supplementary Figure 5B). The overexpression of mtRNAP-3HA was confirmed with Western blot (Supplementary Figure 5B inlet). Importantly, with Blue Native PAGE, we could show, that the overexpressed mtRNAP is able to integrate in the high-molecular weight complex (Supplementary Figure 5C). However, the overexpression had neither an effect on mini- or maxicircle DNA abundance (Supplementary Figure 5D) nor on maxicircle transcription (Supplementary Figure 5E). Next, we overexpressed mtTF1 tagged with a 3xHA tag. Similarly, the overexpressed mtTF1-HA localizes not only to the mitochondrial DNA but is dispersed in the mitochondria (Supplementary Figure 6A). The overexpression did not affect cell growth (Supplementary Figure 6B) and had no effect on mini- and maxicircle DNA and maxicircle transcription (Supplementary Figure 6C and D). We conclude that the overexpression of neither mtRNAP nor mtTF1 impairs cell growth.

## Discussion

This study reports the identification of a novel protein, herein referred to as mitochondrial transcription factor 1 (mtTF1) in the single celled eukaryotic parasite *T. brucei*. We demonstrate that mtTF1 physically and functionally interacts with the mitochondrial RNA polymerase (mtRNAP), and that both proteins are essential for mitochondrial gene expression and the maintenance of one specific component of the mitochondrial genome, the maxicircle DNA.

Functional interrogation via RNA interference revealed a strand-specific effect on steady state level transcript abundance following mtRNAP and mtTF1 depletion. Transcripts derived from genes located on the minor strand exhibited a more pronounced reduction than those on the major strand. One plausible explanation is differential promoter strength, wherein transcription from the major strand promoter is maintained more robustly under limiting conditions during RNAi knockdown. This observation suggests a preferential recruitment of the transcriptional apparatus—comprising at least mtRNAP and mtTF1—to the major strand promoter, consistent with higher promoter activity. A comparable phenomenon has been reported in mammalian systems; in mouse cardiac mitochondria, reduced mtRNAP levels favour transcription initiation at the light strand promoter over the heavy strand promoter (Kühl et al., 2016). This has significant implications since the light strand promotor-initiated transcripts can serve as primers for mitochondrial DNA replication, suggesting a prioritization of the transcription machinery for replication purposes under constrained polymerase availability (Falkenberg et al., 2007; Gustafsson et al., 2016; Kühl et al., 2016). Whether a similar mechanism operates in *T. brucei* remains to be confirmed, as detailed mapping of promoter elements and assessment of their affinities for mtRNAP and mtTF1 remain unknown.

It is still debated whether mitochondrial gene expression is mono- or polycistronic in *T. brucei*. Evidence for monocistronic transcription is based on the occupancy of mtRNAP with individual genes, instead of uniform binding of mtRNAP to the whole mitochondrial genome (Sement et al., 2018). However, earlier studies using Northern blots identified polycistronic transcripts (Feagin et al., 1985; Koslowsky & Yahampath, 1997; Read et al., 1992). Our RNA sequencing and RT-qPCR analyses support the presence of polycistronic (or multicistronic) transcription units. Specifically, we observed that transcripts of the same strand were similarly affected by mtRNAP and mtTF1 knockdown. This coordinated expression pattern suggests that genes may be transcribed as a single RNA molecule from a shared upstream promoter, consistent with polycistronic transcription. The partial protection of major strand steady state transcript levels upon mtRNAP and mtTF1 RNAi further implies the presence of promoters with different strengths.

Additionally, the higher 12S rRNA levels compared to ND4 mRNA levels upon mtRNAP knockdown may reflect its incorporation into ribosomal complexes which protect it from degradation (Gelfand & Attardi, 1981). Intriguingly, we observed a two-fold increase of the 12S rRNA in yL262P cells after ten days of mtRNAP RNAi induction, which might indicate enhanced mitochondrial ribosome biogenesis and thereby protection of 12S rRNA. A similar phenomenon was observed in mitochondrial leucine-rich pentatricopeptide repeat containing (LRPPRC) knockout mice. LRPPRC regulates mitochondrial mRNA stability and its knockout in mice upregulates mitochondrial ribosome biogenesis to compensate for respiratory chain deficiency (Ruzzenente et al., 2012). It remains to be elucidated if yL262P cells, which do not depend on mitochondrial oxidative phosphorylation, show increased mitochondrial ribosome biogenesis upon impairment of mitochondrial gene expression.

We observed a decrease of maxicircle DNA abundance upon mtRNAP and mtTF1 RNAi. Similarly, disruption of mitochondrial transcription factors in other systems has been linked to defects in mtDNA replication. For instance, yeast Mtf1 and human TFAM and TFB2M are essential for mtDNA maintenance and knockout results in decreased mtDNA levels (Ekstrand et al., 2004; Inatomi et al., 2022; Jiang et al., 2011). Furthermore, human TEFM is important for de novo mtDNA synthesis and mTERF may modulate mtDNA replication pausing (Hyvärinen et al., 2007; Matsuda et al., 2025). Therefore, it is not surprising, that mtTF1 knockdown results in decreased maxicircle DNA levels.

In *T. brucei*, the mitochondrial primase Pri1 has previously been implicated in maxicircle DNA replication, while a second mitochondrial primase, Pri2, was proposed to function primarily in the replication of minicircles (Hines & Ray, 2010, 2011). However, our findings that the mtTF1– mtRNAP complex is also essential for maxicircle maintenance prompt a reassessment of these earlier assignments. Maxicircle replication is generally believed to occur within the central region of the kinetoplast DNA (kDNA) disk, whereas minicircle replication is localized to peripheral regions termed antipodal sites (Amodeo et al., 2023; Gluenz et al., 2011). Notably, mtTF1 and mtRNAP are localized within the kDNA disk, in spatial concordance with maxicircle replication, whereas Pri1 localizes to the antipodal sites, in association with other proteins implicated in minicircle replication (Amodeo et al., 2023; Gluenz et al., 2011).

Functional studies further support this spatial-functional distinction. RNAi-mediated knockdown of Pri1 produces a phenotype characterized by extensive kDNA fragmentation, consistent with a primary defect in minicircle replication. A comparable phenotype is observed following depletion of the minicircle-specific DNA polymerase PolIC (Klingbeil et al., 2002). In both cases, the concurrent loss of minicircles and maxicircles, along with structural disintegration of the kDNA network, suggests that maxicircle depletion is likely a secondary consequence of compromised minicircle replication and kDNA network integrity. In contrast, knockdown of mtRNAP or mtTF1 leads to a selective reduction in maxicircle DNA abundance without affecting minicircle levels, indicating a specific and direct role in maxicircle maintenance. These findings strongly support the functional segregation of mitochondrial primases and transcription machinery with respect to distinct components of the kinetoplast genome.

Our findings establish mtTF1 as a key partner of mtRNAP. Together, the two proteins are crucial for mitochondrial transcription and replication in *T. brucei.* While mtRNAP is conserved among different eukaryotic organisms, mtTF1 appears unique to the Kinetoplastea. Given its essential role in cell survival, mtTF1 is a promising candidate for drug development in this group of human and animal pathogens.

## Materials and Methods

### Transgenic cell lines

Transgenic *T. brucei* cell lines were generated using the procyclic form (PCF) strain 29-13 and the bloodstream form (BSF) strain NYsm and yL262P (Dean et al., 2013; Motyka & Englund, 2004; Wirtz et al., 1999). The PCF were grown in exponential phase (up to 1×10^7^ cells/ml) in semi-defined medium 79 (SDM-79, Bioconcept, #9-04V01-M) supplemented with 10% fetal calf serum (FCS) at 27 °C. BSF were grown in exponential phase (up to 1×10^6^ cells/ml) in Hirumi’s modified Iscove’s medium 9 (HMI-9) at 37° C (with 5% CO_2_ and water) (Hirumi & Hirumi, 1989). HMI-9 was supplemented with 10% FCS. For the PCF, antibiotics were added to a final concentration of neomycin (15 µg/ml), hygromycin (25 µg/ml), puromycin (1 µg/ml) and blasticidin (10 µg/ml). For BSF, antibiotics were added to a final concentration of neomycin (2.5 µg/ml), phleomycin (2.5 µg/ml), hygromycin (2.5 µg/ml), puromycin (0.5 µg/ml), blasticidin (5 µg/ml).

Tagging was performed as described by Oberholzer and coworkers (Oberholzer et al., 2006). Briefly, the forward primer was designed to contain the last 100-120 bases of the 3’ end of the target gene, and the reverse primer contains the sequence of the 3’ untranslated region (UTR) after the target gene in reverse complement. Primers are listed in Table 1. Sequences were added to the primers to allow for amplification of a tag (Myc or HA) and a resistance gene (puromycin or blasticidin resistance gene) from the pMO plasmids by PCR. The amplified tag with the resistance gene was cleaned up according to the QIAquick ^®^ PCR Purification Kit (QIAGEN; #28106). This PCR product was used for transfection.

**Table 1:**
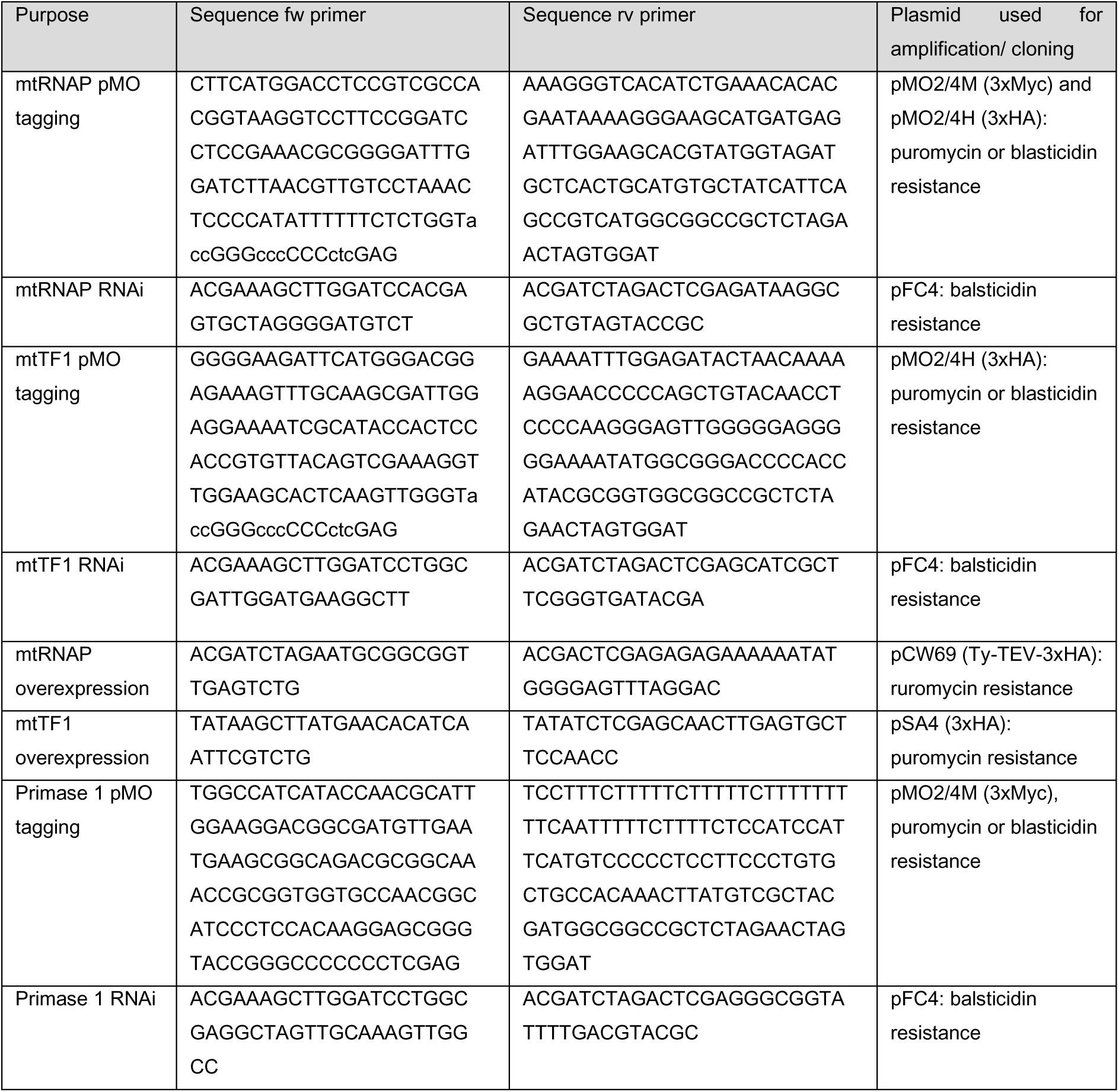
Primer list for PCR. The table includes information about the purpose for which the primers were used, the sequence of the forward and reverse primer as well as the plasmid used for amplification or cloning, respectively.

For the RNA interference (RNAi), a stem-loop construct containing the RNAi sequences in opposite direction was cloned into a modified pLew100 vector (pFC4) which contains a blasticidin resistance cassette (Bochud-Allemann & Schneider, 2002). The construct is inducible with tetracycline, which results in the expression of an RNA hairpin structure that is complementary to the target RNA and results in its degradation via RNAi. The RNAi targets the open reading frame of mtRNAP (position: 575-1045), mtTF1 (position: 98-567) and Primase 1 (position: 40-452). The RNAi target was selected using the RNAi selection tool from Redmond and coworkers (Redmond et al., 2003). Importantly, restriction sites were added to the primers: the forward primer contains HindIII (BioConcept, #R3104L) and BamH (Bioconcept, #R0136L), whereas the reverse primer contains XhoI (BioConcept, #R0146M) and XbaI (BioConcept, #R0145T) restriction sites. Prior to transfection, the plasmid was linearized with Not1 (BioConcept, #R3189M), and cleanup was performed using the QIAquick ^®^ PCR Purification Kit (QIAGEN; #28106).

Transfection was performed with 1×10^8^ PCF and 5×10^7^ BSF trypanosomes, respectively. Cells were harvested (2800 rpm, 5 min, RT) and resuspended in 110 µl transfection buffer containing 10 µg of DNA (Schumann Burkard et al., 2011). Cells were transferred to a transfection cuvette and transfected with the Amaxa Nucleofector II program X-014 for PCF and X-001 for BSF trypanosomes, respectively. Upon transfection, cells were transferred to 10 ml SDM-79 (PCF) and 40 ml of warm HMI-9 (BSF), respectively. To obtain clonal cultures, these pools were diluted 1:10, 1:100 and 1:1000 and distributed in 24-well plates for each of the dilutions. Antibiotics were added 20 h (PCF) and 6 h (BSF) after transfection. Usually, clones were recovered 14 days after transfection for PCF and five days after transfection for BSF. The following PCF cell lines were generated: mtRNAP-3Myc (puromycin) x mtRNAP RNAi (blasticidin), mtTF1-3HA (puromycin) x mtTF1 RNAi (blasticidin), mtRNAP-3HA (puromycin) x mtTF1 RNAi (blasticidin), mtTF1-3HA (puromycin) x mtRNAP RNAi (blasticidin), mtTF1-Myc (blasticidin) x mtRNAP-HA (puromycin). Primase1-3Myc (puromycin) x Primase1 RNAi (blasticidin), mtRNAP-3HA overexpression (puromycin), and mtTF1-3HA overexpression (puromycin). The following BSF cell lines were generated: mtRNAP-3Myc (puromycin) x mtRNAP RNAi (blasticidin) in NYsm and mtRNAP-HA (phleomycin) x mtRNAP-RNAi (blasticidin) in yL262P.

### Maximum-likelihood based phylogeny

We downloaded the protein sequences from TriTryp DB (https://tritrypdb.org/tritrypdb) and used the multiple sequence alignment tool MUSCLE to align the protein sequences for phylogenetic reconstruction (Edgar, 2004). For reconstruction, we used a maximum-likelihood-based phylogeny approach in combination with a Shimodaira-Hasegawa-like branch test (Guindon et al., 2010). The visualisation of the tree was done using TreeDyn (Chevenet et al., 2006).

### Western Blot

Whole-cell lysates were prepared for Western blot as follows: cells were harvested at 4000 rpm for 8 min and washed once with 1 ml 1x phosphate-buffer saline (PBS). Afterwards, cells were resuspended in 1x Laemmli buffer containing 12 mM Tris-HCl pH 6.8, 0.4% SDS, 2% glycerol, 1% β-mercaptoethanol, and 0.002% bromophenol blue in PBS. Denaturation was carried out at 95°C for 5 min. 5×10^6^ cells were loaded per lane onto a 10% SDS-PAGE. The SDS-PAGE was run at 80 V until the samples reached the resolving gel. The voltage was increased to 100 V, and the SDS-PAGE run until the samples reached the end of the gel. Subsequently, proteins were transferred to a nitrocellulose membrane with semi-dry transfer. Two Whatman filter papers were soaked in transfer buffer (250 mM Tris, 192 mM glycine, 20% ethanol, and 0.01% SDS) and used below and on top of the SDS-PAGE and nitrocellulose membrane (Amersham^TM^ Protran^®^ Premium Western-Blotting-Membrane, Nitrocellulose. Cytiva; #10600003). Semi-dry transfer was carried out at 10V with 120 mA (240 mA for two gels) for 1 h at room temperature. Prior to antibody incubation, unspecific binding of antibodies was reduced by blocking the membrane in 5% skim milk powder diluted in PBS containing 0.1% Tween^®^ 20 (Sigma-Aldrich; #93773, PBS-T) at room temperature for 30 min. Primary and HRP conjugated secondary antibodies were diluted in PBS-T containing 5% milk with a dilution of 1:1000 and 1:10000, respectively. Antibodies are given in Table 2. Antibodies The membrane was incubated with antibodies for 1 h at RT for each antibody with washing in between primary and secondary antibody incubation. Washing was repeated three times with PBS-T at room temperature for 5 min per each wash. After a final wash in PBS, the proteins were visualized using the SuperSignal West Femto Maximum Sensitivity Substrate (ThermoFisher Scientific; #34096). Chemiluminescent signal was detected with the Amersham Imager 600 (GE Healthcare) in semi-automated mode.

### Immunofluorescence Assay

For immunofluorescence (IFA), 1×10^6^ *T. brucei* cells were harvested (2800 rpm, 5 min, RT). After washing once with PBS, cells were resuspended in 10 µl PBS and spread on a microscopy slide. Cells were then fixed with 4% paraformaldehyde (PFA) in PBS for 4 min at room temperature, followed by two PBS washes. Cells were permeabilized with 0.2% Triton^TM^ X-100 (Sigma Aldrich, 9036-19-5) for 4 min at room temperature, followed by two additional PBS washes. To reduce nonspecific antibody binding, cells were blocked with 4% bovine serum albumin in PBS (BSA, Sigma-Aldrich, #A2153) for 30 min at RT. Afterwards, cells were incubated with primary antibodies (Table 2) diluted in 4% BSA for 1 h at room temperature. The cells were then washed three times for 5 min with PBS and subsequently incubated with secondary antibodies (Table 2) at room temperature for 1 h followed by three more washes with PBS. The samples were mounted with 10 μl ProLong^TM^ Gold Antifade Mountant with DNA stain DAPI (ThermoFisher, #P36931). Images were acquired with a widefield Leica DM5500 B microscope with a 100x oil immersion phase contrast objective. LAS AF software (Leica) and ImageJ were used for image analysis.

**Table 2:** Antibodies used for Western blot and immunofluorescence assay. The antibody dilution factor and provider are indicated.

| Target | Dilution | Provider and catalogue number |
| --- | --- | --- |
| ATOM40 rabbit | 1:1000 | Kind gift from André Schneider (Mani et al., 2015) |
| EF1α mouse | 1:1000 | Santa Cruz, #SC-21758 |
| COXIV | 1:1000 | Kind gift from André Schneider (Niemann et al., 2013) |
| mtHSP70 | 1:1000 | Kind gift from Luise Krauth-Siegel |
| HA rat | 1:1000 | Roche, #11867423001 |
| Myc mouse | 1:1000 | Sigma Aldrich, #M4439 |
| Anti-mouse HRP | 1:10000 | Dako, #P0260 |
| Anti-rat HRP | 1:10000 | Dako, #P0450 |
| Anti-rabbit HRP | 1:10000 | Dako, #P2017 |
| Anti-mouse 488 | 1:1000 | ThermoFisher, #A11001 |
| Anti-rat 594 | 1:1000 | ThermoFisher, #A11007 |

### Digitonin extraction

Mitochondria were enriched using digitonin extraction as described by Schneider and coworkers (Schneider et al., 2007). Briefly, 5x10^7^ cells were harvested by centrifugation (2000 rpm for 7 min at RT) and washed once with 1 ml PBS. The pellet was resuspended in 250 µl 1x SoTE buffer (0.6 M sorbitol, 20 mM Tris-HCl pH 7.5, 2 mM EDTA with pH 8) and transferred to a 1.5 ml tube. To selectively permeabilize the plasma membrane, 250 µl 1x SoTE containing 0.05% digitonin (BIOSYNTH; #D3203) was gently added to obtain a final digitonin concentration of 0.025%. Cells were incubated on ice for 5 min. After centrifugation at 8000 rpm, 4°C, for 5 min, 400 µl of the supernatant (cytosolic fraction) was transferred to a new tube, and 100 µl 5x Laemmli buffer was added. The pellet, containing the mitochondria-enriched fraction, was resuspended in 400 µl 1x SoTE, followed by the addition of 100 µl Laemmli buffer. All samples were boiled at 95°C for 5 min. For Western blot analysis, 1.5×10^6^ cells were loaded per lane for the whole-cell fraction, cytosolic supernatant and mitochondria-enriched pellet.

### Co-immunoprecipitation (Co-IP)

Mitochondria were isolated using digitonin extraction, as described above. Mitochondria were lysed in lysis buffer (20 mM Tris HCl pH 7.4, 100 mM NaCl, 0.1 mM EDTA, 25 mM KCl) supplemented with 1.5% digitonin and EDTA-free Complete^™^ ULTRA Tablets EASYpack Protease Inhibitor Cocktail (Sigma Aldrich, # 5892791001) at five times the recommended concentration. Samples were incubated on ice for 15 min. Afterwards, samples were centrifuged at 14000 rpm, 4°C, for 15 min. The supernatant, containing the soluble mitochondrial proteins, was used as input for the immunoprecipitation. The supernatant was incubated with 25 µl prewashed Anti-HA Magnetic Beads (Sigma Aldrich, #SAE0197) for PCF and EZview^TM^ Red Anti-c-Myc Affinity Gel beads (Sigma Aldrich, #E6654) for BSF. Additionally, EZview^TM^ Red Anti-HA Affinity Gel beads (Sigma Aldrich, #E6779) were used for the reciprocal IP experiments. Binding was performed at 4°C for 1 h. Beads were pelleted using either a magnetic holder or centrifugation at maximum speed, for 10 seconds. The supernatant (flowthrough) containing unbound proteins, was transferred to a new tube. Beads were subsequently washed five times with 500 µl wash buffer (20 mM Tris HCl pH 7.4, 100 mM NaCl, 0.1 mM EDTA, 25 mM KCl) containing 0.1% digitonin and EDTA-free complete^™^ ULTRA Tablets EASYpack Protease Inhibitor Cocktail (Sigma Aldrich, # 5892791001) at five times the recommended concentration. For subsequent mass spectrometry, three additional washes were performed using digitonin-free wash buffer. Data and further detail of the mass spectrometry protocol are publicly available on ProteomeXchange Consortium. For western blot analysis, samples were prepared and loaded as described above.

### RNA Isolation

A total of 1×10^8^ cells were harvested by centrifugation at 2500 rpm for 8 min at room temperature. The pellet was washed with 1 ml PBS, transferred to a 1.5 ml tube and centrifuged again at 2500 rpm for 8 min at room temperature. After removing the supernatant, the cell pellet was resuspended in 1 ml TRIzol (Tri-Reagent, Merck, #9424). Subsequently, 200 µl of chloroform was added and the samples were vortexed vigorously. The mixture was then centrifuged at 12000 rpm for 15 min at 4°C. The upper aqueous phase (400 µl) was carefully transferred to a new tube containing 500 µl of ice-cold isopropanol, ensuring that the interphase was not disturbed. The tube was inverted several times to mix the solution. RNA was precipitated by centrifugation at 12000 rmp for 15 min at 4°C. Subsequently, the supernatant was removed and discarded. The RNA pellet was washed twice with 1 ml 70% ethanol followed by centrifugation at 7500 rpm for 5 min at 4°C. The ethanol was removed, and the tube was inverted and left open at room temperature to air-dry for at least 5 min. Finally, the RNA pellet was resuspended in 50 µl RNase-free water and incubated at 55°C for 5 min on a heating block. RNA concentration was determined using a Nanodrop spectrophotometer and RNA integrity was assessed by running 1 µg of RNA on a 1% agarose gel for 40 min at 100 V. RNA was stored at -80°C or used directly for DNase treatment and cDNA synthesis.

### Northern blot analysis for small mitochondrial RNAs

**1. RNA Isolation from digitonin-extracted mitochondria:** Mitochondria-enriched pellets were obtained as described above with digitonin extraction. RNA was isolated from these pellets following the RNA isolation protocol outlined above. **2. Polyacrylamide Gel Electrophoresis (PAGE) and Northern blot transfer**: Denaturing PAGE was performed using 10% urea-TBE (Tris borate EDTA) gels (Bio-Rad, #4566033). A total of 10 µg of RNA was heated at 95°C for 60 sec and mixed with 2x RNA loading dye (95% formamide, 0.025% bromophenol blue, 0.025% xylene cyanol). Electrophoresis was conducted in 1x TBE running buffer at a constant 200 V for 90 min. Following electrophoresis, gels were stained in 0.5x TBE buffer containing RedSafe for 5 min, and gel images were captured before transfer. RNA was transferred onto a positively charged nylon membrane (Roche, #11417240001) via semi-dry electroblotting at 300 mA for 1 h in 0.5x TBE buffer. Subsequently, the membrane was crosslinked with UV light at 0.12 J. **3. Probe labelling and hybridization:** Oligonucleotide probes (20-30 nt) complementary to the target RNA were 5’-end labelled using T4 polynucleotide kinase (PNK, NEB, #M0201S). Based on the RNAseq data, we chose an abundant gRNA and ordered short oligonucleotides complementary to the gRNA of ND7 encoded on mO_236:667:716 (Cooper et al., 2022). The oligonucleotide has the sequence 5’-TATTATCATTCATACTCACTGTACTATAAGTTATCTGCATCGTATTATAT-3’. The labelling reaction (total volume 10 µl) included 1.2 µl oligos (10 pmol/µl), 1 µl 10x PNK buffer, 5 µl H2O, 2 µl ɣ-32P-ATP, and 0.8 µl T4 PNK (10,000 U/ml). The reaction was performed at 37°C for 30 min, followed by heat inactivation at 95°C for 3 min. Membranes were prehybridized in 15-20 ml hybridization buffer at 42°C before adding the radiolabelled probe. Hybridization was carried out at 42°C for 18 h. **4. Washing and Imaging** 20 x SSC wash buffer was prepared: 3 M NaCl and 300 mM Tri Sodium citrate were dissolved in water and the pH was adjusted with HCl to 7. Following hybridization, membranes were washed first in prewarmed 2x SSC (0.1% SDS) for 10 min at 42°C followed by 0.5x SSC (0.1% SDS) for 10 min at 42°C and lastly by briefly rinsing the membrane in water at RT. The membranes were air-dried, wrapped in plastic wrap, and exposed to a phosphor screen for 72 h. RNA signals were visualized using a Storm PhosphoImager (Amersham Bioscience).

### DNA Isolation

5×10^7^ cells were harvested by centrifugation at 2000 rpm for 5 min. After removing the supernatant, the cells were washed in 1 ml PBS and transferred to a 1.5 ml tube. Following centrifugation at 3000 rpm for 5 min, the pellet was resuspended in 100 µl DNA QuickExtract DNA Extraction Solution (LubioScience, #QE0905T) and vortexed for 15s. Samples were then incubated at 65°C for 6 min, vortexed again for 15s, and incubated at 98°C for 2 min. The DNA concentration was measured using a Nanodrop spectrophotometer. Here, the 260/280 ratio was typically below 1. The reproducibility of this method was verified by comparing mtRNAP and Primase 1 RNAi data with published literature (Grams et al., 2002; Hines & Ray, 2010). In addition to being extremely fast, the method was verified to be reproducible. DNA concentration was adjusted to 100 ng/µl and stored at -20°C. For qPCR, a DNA working concentration of 4 ng/µl was used.

### RNA-sequencing, qPCR and RT-qPCR

For DNase treatment, a total amount of 10 µg RNA was used. In a final volume of 100 µl, 10 µl DNase buffer (NEB, #B0303S), 1 µl DNase (NEB, #M0303S) and 10 µg RNA were incubated at 37°C for 15 min. Afterwards, the RNeasy MinElute Cleanup kit (QIAGEN, #74204) was used to purify the RNA prior to cDNA synthesis. It is crucial to obtain pure RNA for reproducibility of RT-qPCR results. After cleanup, the RNA was sent to FASTERIS for RNA sequencing (one replicate for mtRNAP and mtTF1 RNAi (each -Tet and +Tet [2d]). The RNA sequencing data and further information about the RNA sequencing protocol and analysis is publicly available at the Gene Expression Omnibus.

For RT-qPCR, the iScript gRNA Clear CDNA synthesis Kit (Biorad, #1725034) was used to synthesis cDNA according to the manufacturer’s protocol. Briefly, 1 µg RNA was used and adjusted to a final volume of 40 µl with water. Subsequently, 10 µl of 5x iScript Super Mix was added, either containing reverse transcriptase or lacking it for the negative control. The cDNA synthesis was performed in a thermocycler using the following program: 25°C for 5 min, 46°C for 20 min, 95°C for 1 min and hold at 4°C. cDNA was stored at -20°C. The cDNA was then diluted 1:10 with water and directly used for RT-qPCR.

All primers used for qPCR and RT-qPCR are listed in Table 3. Primers for detecting minicircles were previously published by Gluenz and coworkers (Gluenz et al., 2011), while primers for ND4 were previously published by Li and coworkers (Li et al., 2011). GAPDH and actin primer were previously published by Brenndörfer and Boshart (Brenndörfer & Boshart, 2010). Primers for the detection of β-tubulin, 12S rRNA, COI, ND1 and Primase 1 were designed using the IDT PrimerQuest^TM^ Tool (https://eu.idtdna.com/PrimerQuest/Home/Index). For each target, three cycle threshold (Cq) values were averaged (pipetting triplicates). Each qPCR analysis was performed in biological triplicates. The comparative Cq method (2^(−ΔΔCq)) was used. Briefly, raw Cq values of target genes were normalized to Cq values of the housekeeping genes (GAPDH or tubulin). Importantly, GAPDH was only used as a housekeeping gene for qPCR but not RT-qPCR, as its mRNA levels increased upon mtRNAP and mtTF1 RNAi. To ensure the validity of tubulin as a housekeeping gene, actin was always run in parallel for RT-qPCR and showed stable expression across conditions. Final Cq values were then further normalized to the uninduced control group for better visualization and are presented as fold change.

**Table 3:**
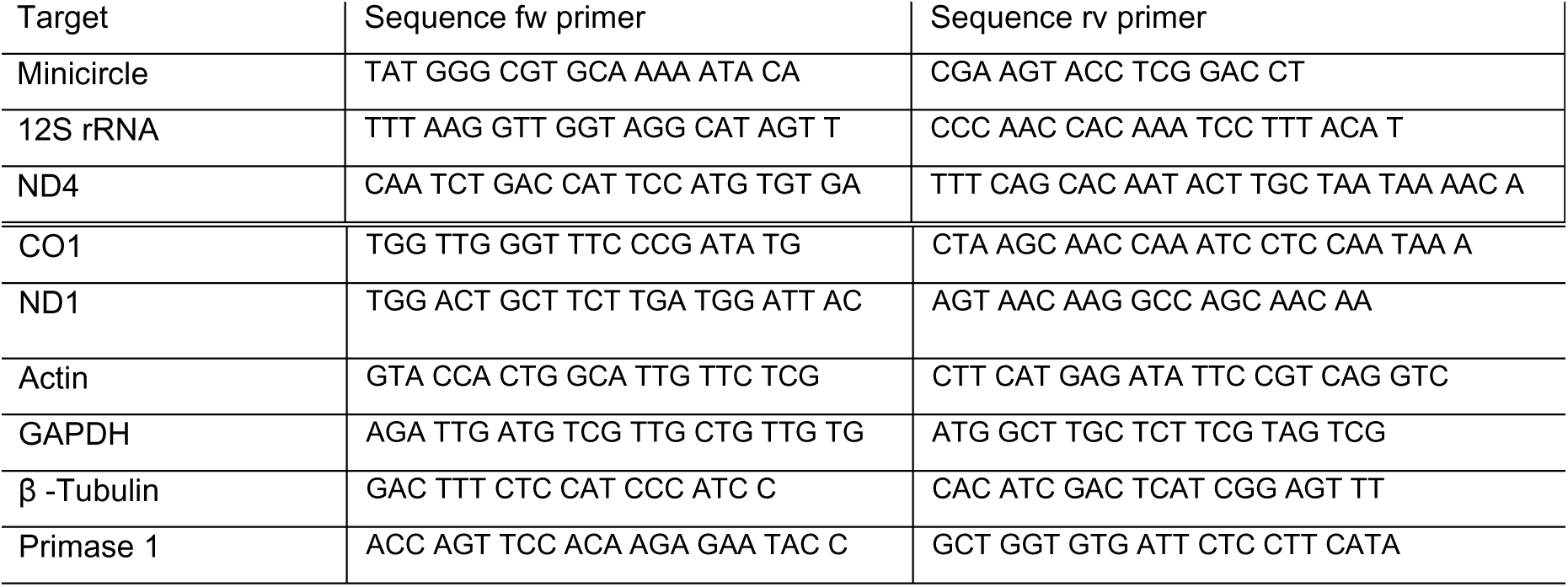
Primers used for qPCR and RT-qPCR. Given are the forward (fw) and reverse (rv) primer for each target detected with qPCR or RT-qPCR.

The qPCR and RT-qPCR were performed as follows: First, primer mixes were prepared for each target. Primers (stock concentration of 100 µM) were diluted to a final concentration of 1.5 µM by adding 15 µl of each primer into 970 µl of water. Subsequently, the primer mix was combined with the Sso Advanced Universal SYBR^®^ Green Supermix (Bio-Rad, #1725271) in a 3:5 ratio. For the RT-qPCR and qPCR, 2 µl of the cDNA (diluted 1:10 after cDNA synthesis) and DNA (at a concentration of 4ng/µl) were pipetted into a 96-well plate. Then, 8 µl of the SYBR^®^ Green Supermix containing the primer mix was added, and the plate was sealed. The thermocycler (Bio-Rad C1000 Touch PCR/Thermal Cycler) program starts at 95°C for 5 min. DNA is amplified during 40 cycles, with each cycle consisting of denaturation at 95°C for 10s and annealing and elongation at 60°C for 15s. Fluorescent intensity was measured after each cycle. During the final step, melting curves were obtained by gradual increasing the temperature from 60°C to 95°C.

### Blue Native PAGE

Mitochondria were isolated from 1×10^8^ cells as described above using digitonin extraction. The mitochondria were then lysed with digitonin solubilization buffer (20mM Tris-HCl pH 7.4, 50 mM NaCl, 5% Glycerol, 0.05mM EDTA, 1 mM PMSF) containing 1% digitonin. Lysis was carried out on ice for 15 min. Subsequently, samples were centrifuges at 14000 rpm, 4°C for 15 min. The supernatant was then used for the Blue Native PAGE, which was performed as previously described (Hoffmann et al., 2018). A total of 5x10^7^ cells were loaded per lane on a native gel with a gradient of 4-13%.

### Pre-gelation staining expansion microscopy (PS-ExM)

The pre-gelation staining expansion microscopy was performed according to Atchou and coworkers (Atchou et al., 2023). Briefly, *T. brucei* cells were stained as described above for regular IFA. The only difference is that 100 µl of washed cells (1×10^6^) was settled on poly-D Lysin functionalized 12 mm coverslips for 15 min and directly fixed with 100 µl 4% of PFA for 4 min (PFA is thereby diluted to a final concentration of 2%). After permeabilization and antibody incubation, cells were incubated in PBS containing 0.7% formaldehyde (FA, 36.5–38%, SIGMA, #F8775) and 1% acrylamide (AA, 40%, SIGMA, #A4058) for functionalization. AA will later allow to link the cells to the hydrogel. Subsequently, samples are embedded into the hydrogel: the gelation solution was prepared freshly with 0.5% ammonium persulfate (APS, 17874, ThermoFisher) and 0.5% tetramethylethylenediamine (TEMED, ThermoFisher, #17919) added to the monomer solution containing 19% sodium acrylate (SA, 97–99%, SIGMA, #408220), 10% acrylamide (AA, 40%, SIGMA, #A4058) and 0.1% N,N’-methylenebisacrylamide (BIS, 2%, SIGMA, #M1533). For gelation, 35 µl of the gelation solution was used per coverslip, and the samples were incubated on ice for 5 min, followed by incubation at 37°C for 30 min. The gels were denatured in 1 ml denaturation buffer (200 mM sodium dodecyl sulfate SDS, 200 mM NaCl and 50 mM Tris in deionized water, pH 9). Ater gently shaking the gels at room temperature for 15 min, the gels detached from the coverslip. Gels were transferred to 1.5 ml tubes and denatured in 1 ml of denaturation buffer at 95°C for 30 min. After a brief incubation of the gels in PBS, the DNA was stained with 5 µg/ml 4′,6-diamidino-2-phenylindole (DAPI, SIGMA, #D9542-5MG) diluted in 2% bovine serum albumin (BSA). DAPI staining was performed at 37°C with gentle agitation for 1 h. Finally, the hydrogels were expanded in deionized water for 1 h. After the final expansion, the gels were cut and mounted on poly-D-lysine functionalized 35 mm glass-bottom dishes (Cellvis, #D35-20-1.5-N. 35 mm glass bottom dish with 20 mm micro-well #1.5 cover glass). Images were acquired with a 60x oil objective (NA= 1.4) on a NIKON Ti 2 CREST V3 equipped with a Hamamatsu Flash 4.0 camera and a Celesta light engine in widefield mode. Images were acquired with a z-step size of 300 nm and pixel size of 108 nm. ImageJ was used for analysis. The Pearson correlation coefficient was measured with the ImageJ plugin JaCoP by selecting only the kDNA region with the rectangular selection tool (n = 10). To calculate the average Pearson correlation coefficient (r), the values were transformed using Fisher’s transformation z = artanh(r), which also allowed to determine the standard error of means (SEM). The obtained average and SEM were then back transformed with the formula r = tanh(z).

## Acknowledgment

The authors acknowledge the lab members of the Ochsenreiter lab for fruitful discussions. We acknowledge Enrico Santini for valuable input. We thank Hauke Hillen and Lea Trost for helpful discussions and reading the manuscript. Furthermore, we acknowledge Prof. André Schneider for discussions and providing the ATOM40 antibody. We thank Robin Eberle, Elena Wettstein, Melanie Steinmann, Ado Crnovrsanin, Roman Engimann and Sandro Käser for help with experiments. The authors gratefully acknowledge Amanda Chantziou, Kodzo Atchou and Christin Berger for their assistance in revising the manuscript. Microscopy was performed on equipment supported by the Microscopy Imaging Centre (MIC) of the University of Bern, Switzerland. We thank the team of the Proteomics and Mass Spectrometry Core Facility (PMSCF) at the Department for BioMedical Research (DBMR) of the University of Bern, Switzerland for their valuable service. The Ochsenreiter lab is supported by grants from the SNSF (207525), the Uniscientia and Novartis foundation.

## Author contributions

BMB, HM, MG and TO conceived and designed the experiments. BMB, HM and MG performed the experiments and analysed the date. B.M.B ,M.G and TO wrote the paper.

---

**Supplementary Figure 1:**
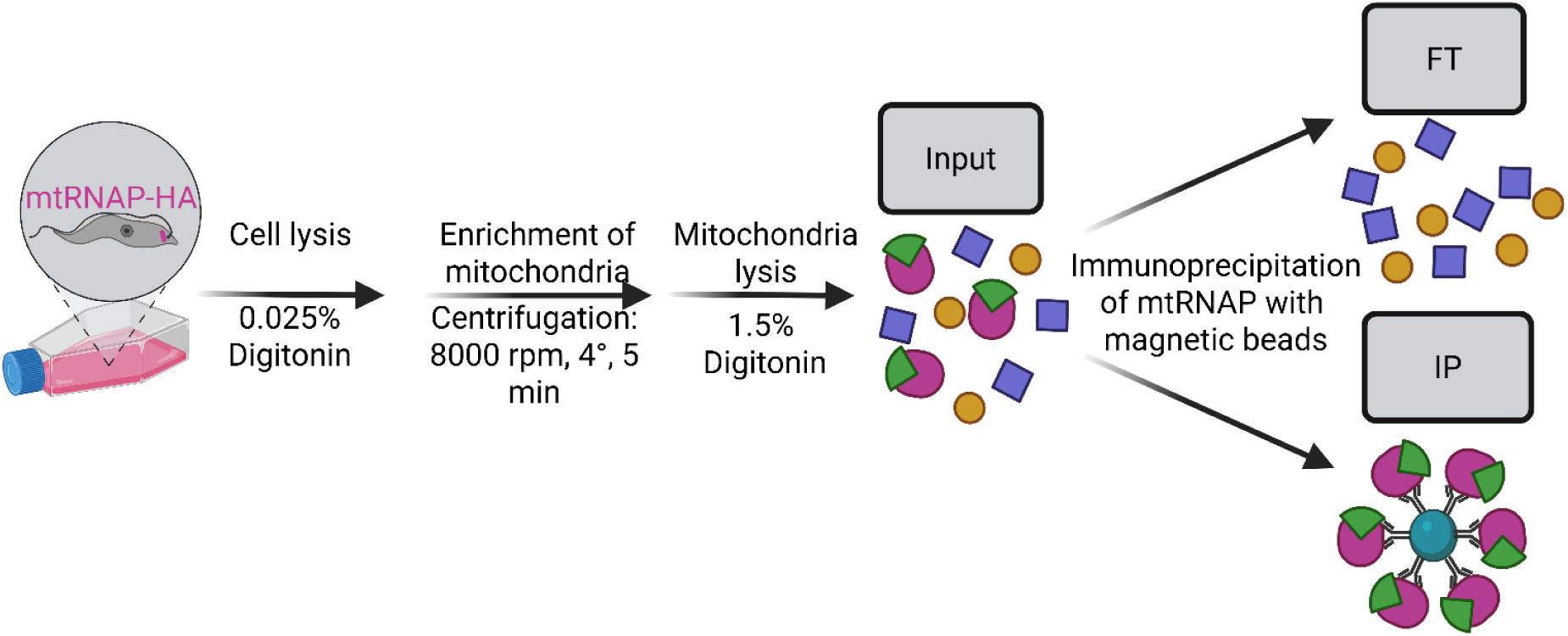
Workflow of the immunoprecipitation of the HA-tagged mtRNAP-HA to identify interaction partners. Cells with C-terminally HA-tagged mtRNAP were lysed with a low percentage of digitonin, and mitochondria were enriched by centrifugation. Following mitochondrial lysis, the proteins (Input) were incubated with magnetic anti-HA beads for 1 h at 4°C. The immunoprecipitation (IP) fraction contains the mtRNAP and potential interaction partners, while other mitochondrial proteins are in the flowthrough (FT). Figure created with BioRender.com.

**Supplementary Figure 2:**
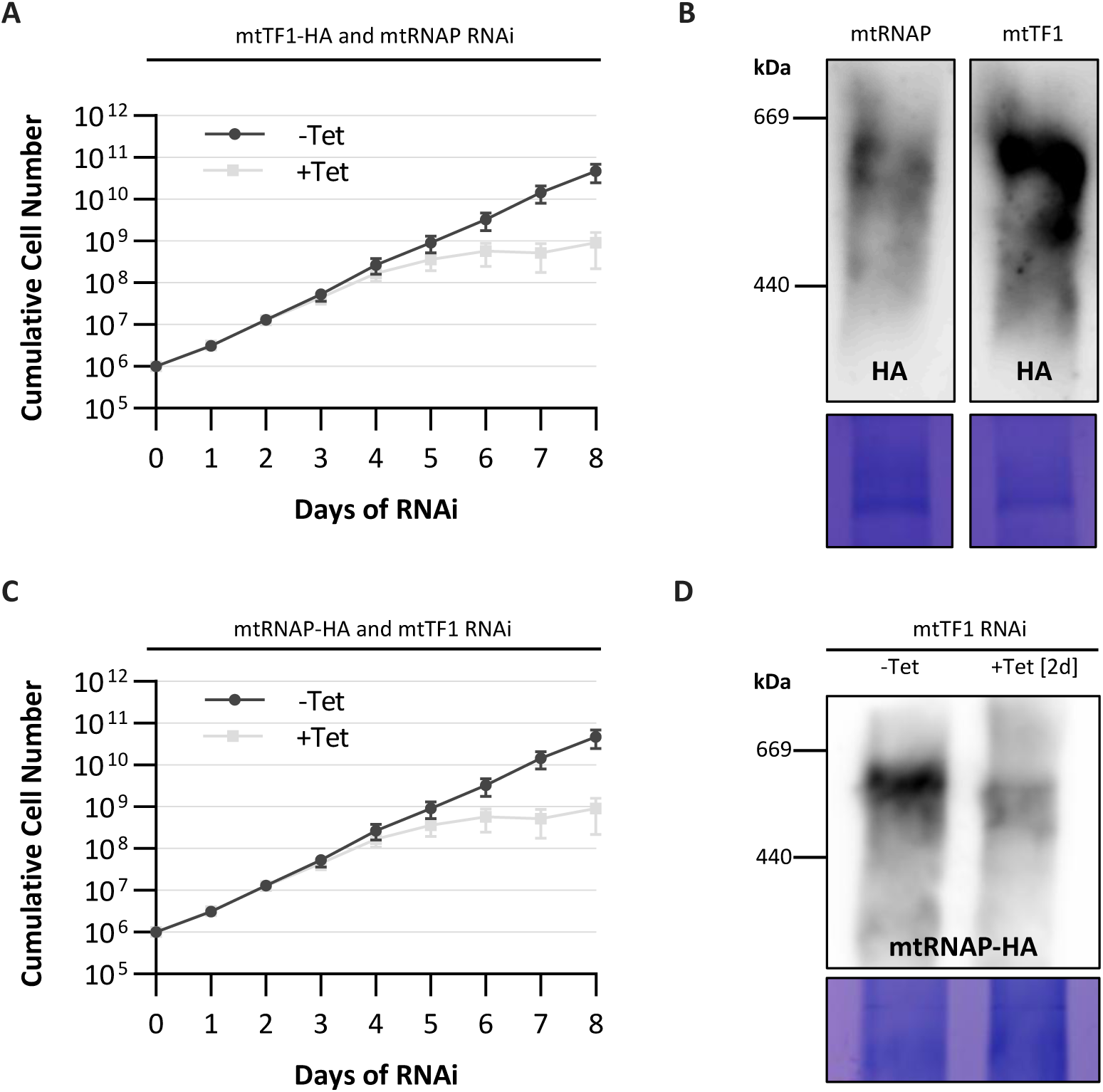
Additional data for mtTF1-HA x mtRNAP RNAi and mtRNAP-HA x mtTF1 RNAi. **(A)** Blue Native PAGE analysis of the mtRNAP and mtTF1 high molecular weight complex. The C-terminally HA-tagged mtRNAP and mtTF1 were detected with antibodies. Coomassie stainings below the Blue Native PAGEs serve as loading control. **(B)** Growth curve of non-induced control cells (-Tet) and mtRNAP RNAi-induced cells (+Tet). Cells express a C-terminally HA-tagged mtTF1. **(C)** Growth curve of non-induced control cells (-Tet) and mtTF1 RNAi-induced cells (+Tet). Cells express a C-terminally HA-tagged mtRNAP. **(D)** Blue Native PAGE analysis of non-induced control cells (-Tet) and mtTF1 RNAi-induced cells (+Tet [2d]). The C-terminally HA-tagged mtRNAP was detected with antibodies. Coomassie staining below the Blue Native PAGE serves as loading control. n = 3 for B and C.

**Supplementary Figure 3:**
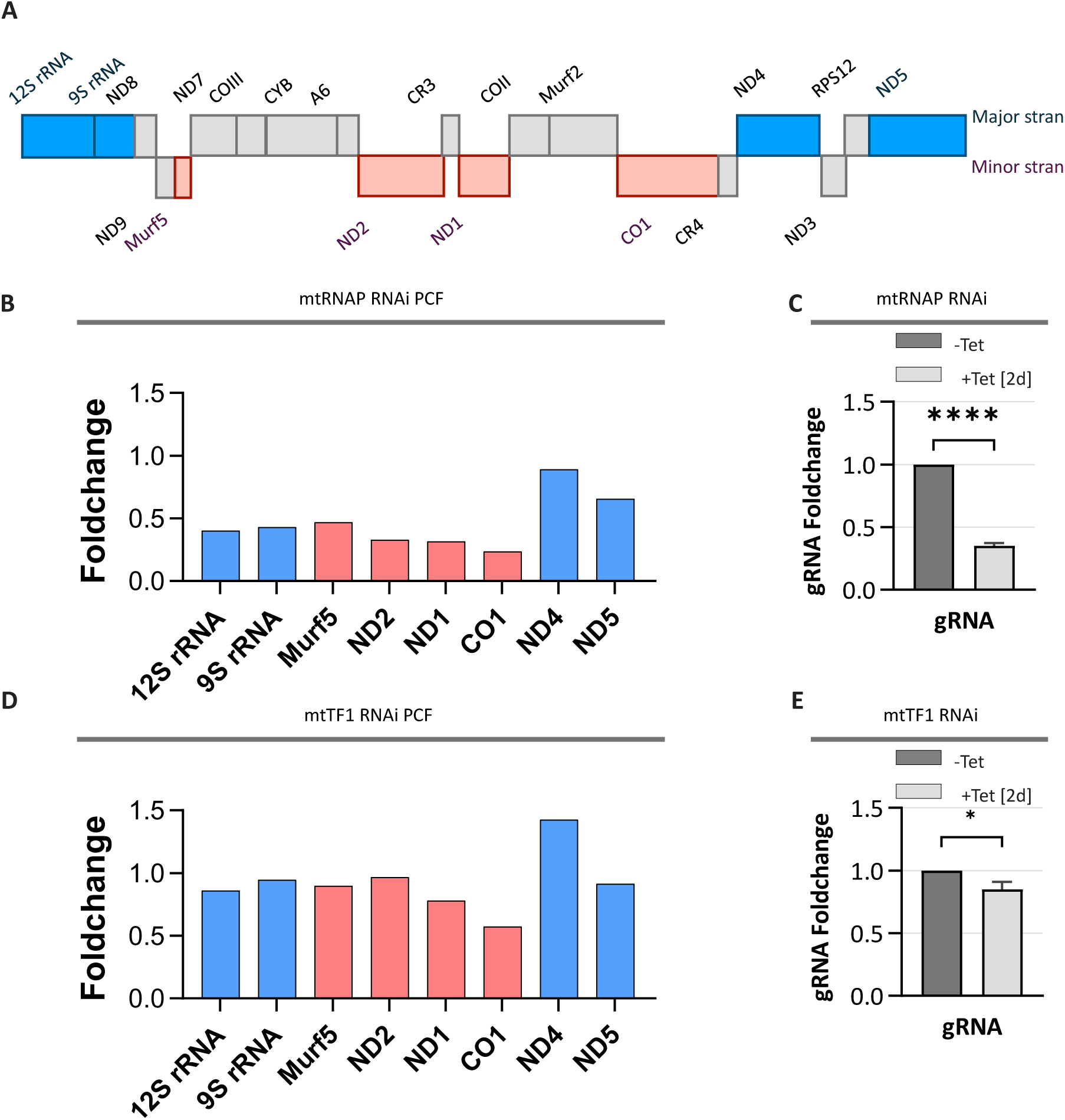
Data analysis of RNA sequencing upon mtRNAP and mtTF1 knockdown. **(A)** Overview of never-edited, maxicircle-encoded transcripts on the major (blue) and minor (red) strand. **(B and D)** Maxicircle transcript foldchange upon mtRNAP and mtTF1 RNAi (+Tet[2d]) shown relative to non-induced control cells. Maxicircle major strand transcript are shown in blue, while minor strand transcripts are shown in red. n = 1. **(C and E)** gRNA foldchange upon mtRNAP and mtTF1 RNAi. Data of gRNA is shown as mean +/- SEM. Different gRNAs with a coverage ≥ 5. have been combined for the analysis.

**Supplementary Figure 4:**
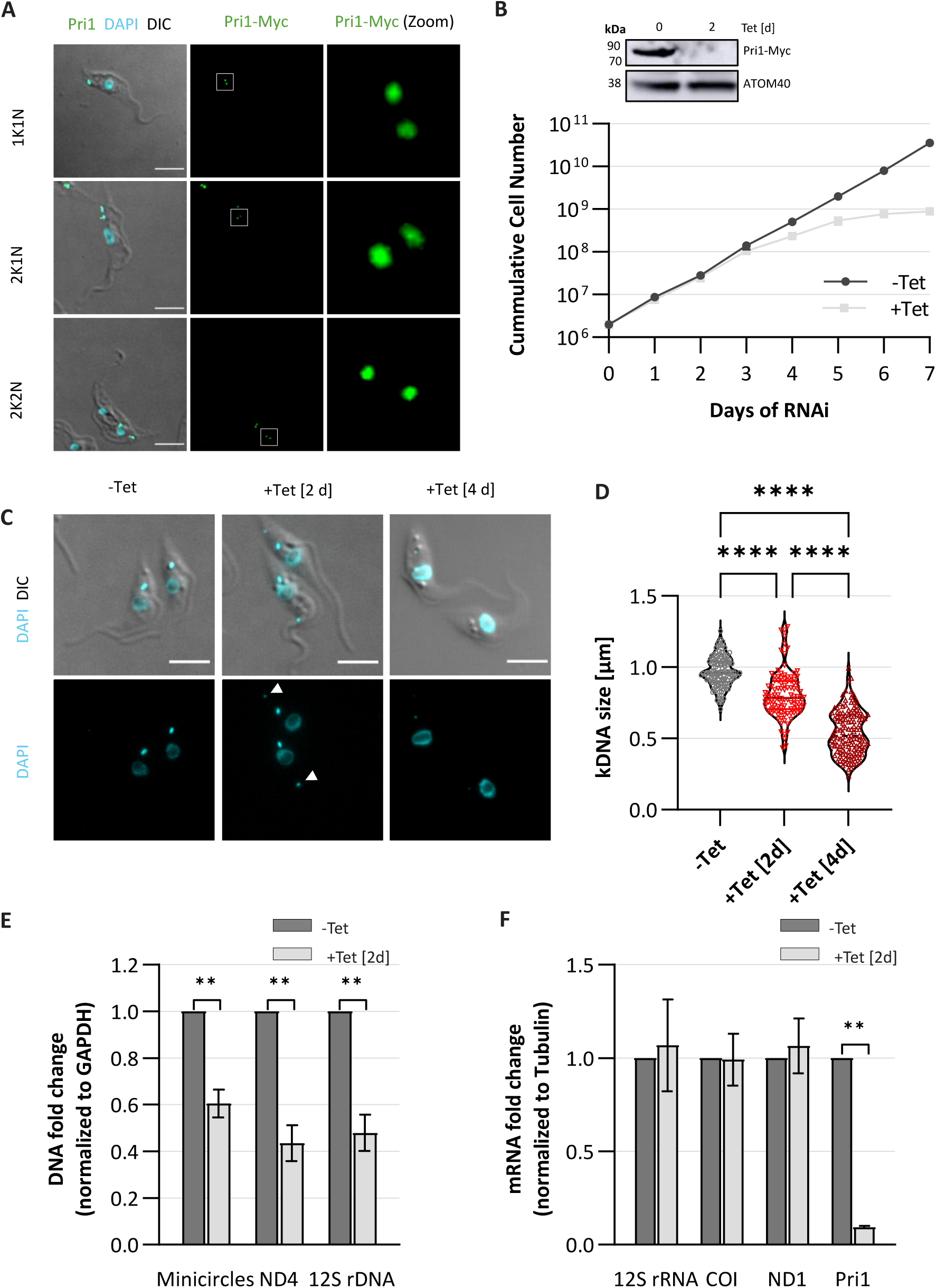
Primase 1 localizes to the antipodal sites flanking the kDNA throughout the cell cycle and is essential for cell viability and kDNA replication in the PCF. **(A)** Immunofluorescence images of cells harbouring a C-terminally Myc-tagged Primase 1 (Pri1, green) together with DAPI staining (DNA, blue). Cells are visualized with DIC (differential interference contrast microscopy). Images show different cell cycle stages. N = nucleus, k = kDNA. **(B)** Growth curve and Western blot analysis of non-induced (-Tet) and Pri1 RNAi-induced (+Tet) cells. ATOM40 served as a loading control. **(C)** Microscopy images showing DAPI staining (DNA, cyan) of non-induced control cells (-Tet) and Pri1 RNAi-induced cells (+Tet [2 and 4d]). Cells are visualized with DIC (differential interference contrast microscopy). **(D)** kDNA size quantification in control cells (-Tet, grey) and Pri1 RNAi-induced cells (+Tet [2d] in red and +Tet [4d] in dark red). **(E and F)** qPCR and RT-qPCR analysis in non-induced control cells (-Tet, dark grey) and upon Pri1 RNAi (+Tet [2d], light grey). Scale bars = 5 µm. Data are shown as mean +/- standard deviation (SD). n = 3 for B, E and F. n = 100 for D.

**Supplementary Figure 5:**
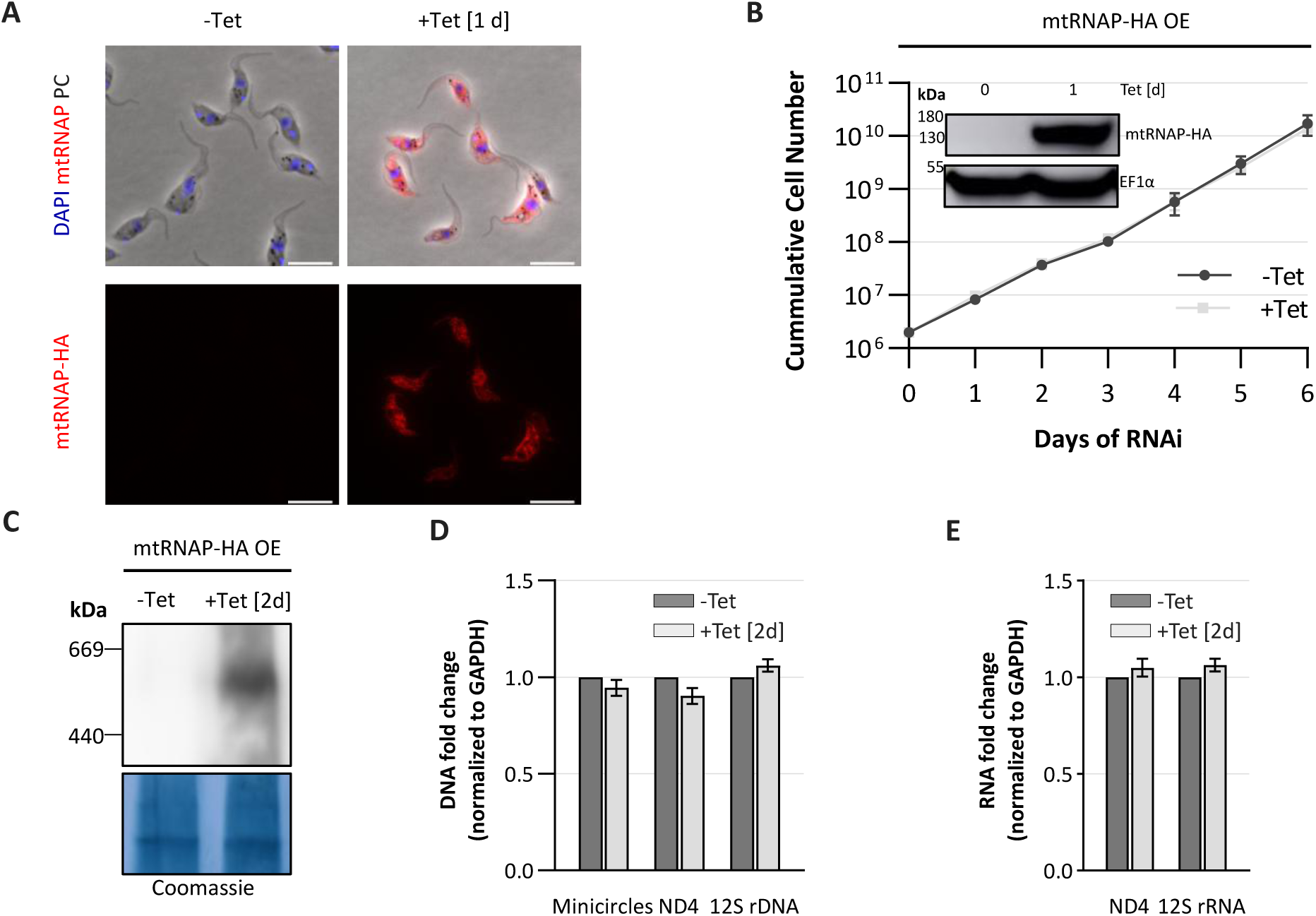
mtRNAP overexpression does not impair cell growth. **(A)** Immunofluorescence images of cells overexpressing mtRNAP-HA (red) upon tetracycline induction together with DAPI staining (DNA, blue). Cells are visualized with phase contrast (PH). **(B)** Growth curve of non-induced control cells (-Tet, dark grey) and inducible mtRNAP-Myc overexpressing cells (+Tet, light grey). Inset: Western blot of non-induced (-Tet) and mtRNAP-HA overexpressing cells (+Tet [1d]). **(C)** Blue Native PAGE probed for HA to detect overexpressed mtRNAP in non-induced (-Tet) and mtRNAP-overexpressing cells (+Tet). **(D and E)** qPCR and RT-qPCR analysis in control (-Tet, dark grey) and mtRNAP-overexpressing cells (+Tet [2d], light grey). Scale bars = 10 µm. Data are shown as mean +/- standard deviation (SD). n = 3 for B, D and E.

**Supplementary Figure 6:**
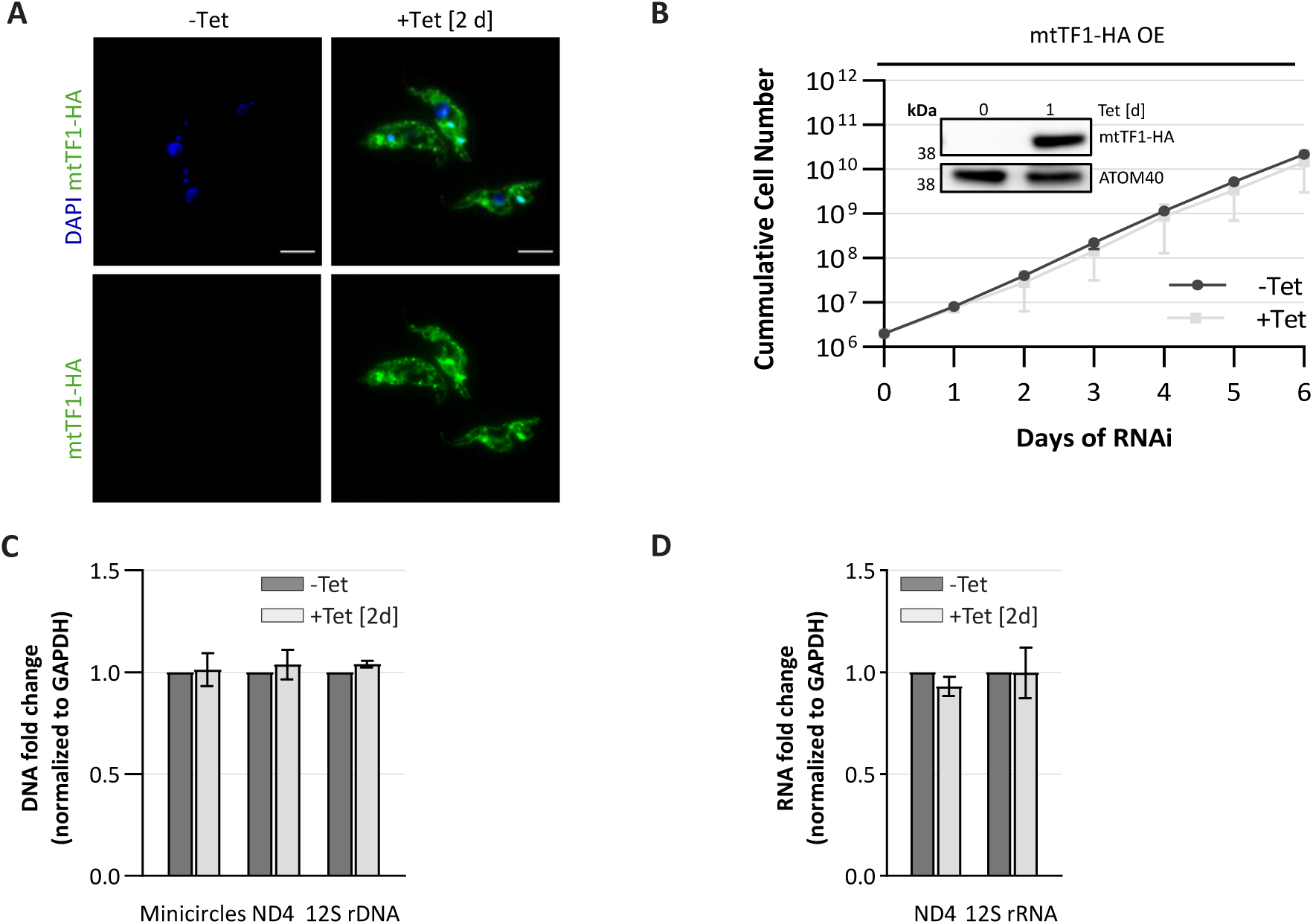
mtTF1 overexpression does not impair cell growth. **(A)** Immunofluorescence images of cells overexpression mtTF1-HA (green) upon tetracycline induction together with DAPI staining (DNA, blue). **(B)** Growth curve of non-induced control cells (-Tet, dark grey) and inducible mtTF1-HA overexpressing cells (+Tet, light grey). Inset: Western blot probed for HA to detect the expression of mtTF1 in non-induced (-Tet) and mtTF1-overexpressing cells (+Tet [1d]). **(C and D)** qPCR and RT-qPCR analysis in non-induced control cells (-Tet, dark grey) and mtTF1-overexpressing cells (+Tet [2d], light grey). Scale bars = 10 µm. Data are shown as mean +/- standard deviation (SD). n = 3 for B, C and D.

